# GlycoForge generates realistic glycomics data under known ground truth for rigorous method benchmarking

**DOI:** 10.64898/2026.02.20.707134

**Authors:** Siyu Hu, Daniel Bojar

**Affiliations:** Department of Chemistry and Molecular Biology, University of Gothenburg, 41390 Gothenburg, Sweden; Wallenberg Centre for Molecular and Translational Medicine, University of Gothenburg, 41390 Gothenburg, Sweden

## Abstract

Quantifying all complex carbohydrates in a sample produces glycomics data, which constitutes compositional data and is stymied by biosynthetic dependencies between glycans, requiring dedicated analytic workflows. Properly assessing such methods frequently requires simulated data with known ground truths and injectable effects. However, simulating glycomics data, especially with control over effects and biases, is still unsolved. Here, we present GlycoForge, a feature-complete solution for simulating comparative glycomics data. GlycoForge supports simulating fully synthetic glycomics data and templated simulations based on real-world data, with specified motif-level effects, based on Gaussian copulas and estimated covariances. We further support injection of batch effects, both mean and variance shifts, via center-log ratio transformations to maintain compositional closure, and realistic missing data simulation. We showcase the utility of GlycoForge by evaluating batch effect correction algorithms for glycomics data, with automated guidelines for when to use such methods on real-world data. GlycoForge is available as an open-access Python package at https://github.com/BojarLab/GlycoForge.

## Introduction

Glycans, the complex carbohydrates decorating proteins and lipids, encode information essential for cellular recognition, immune responses, and disease progression^1–3^. Yet, unlike transcriptomics and proteomics, which benefit from mature computational infrastructures to analyze resulting data downstream, glycomics remains analytically challenging due to fundamental properties of the data itself, such as compositionality and biosynthetic dependencies^4,5^. Glycomics data, when presented as relative abundances, are inherently compositional, existing as proportions that sum to unity within each sample^5^, and an increase in one glycan must mean an artificial “decrease” in all other glycans, to maintain the sum of 100 percent. This creates statistical dependencies that violate assumptions underlying conventional differential expression workflows^6,7^, leading to inflated false discovery rates. While compositionality is to some extent a data representation problem, the biosynthetic pathways producing glycans introduce additional correlations and constraints^8–10^, as enzymes compete for common substrates and precursor structures branch into related products. Therefore, a loss of terminal LacNAc motifs and gain of terminal sialylation (capping those very same LacNAc motifs) might be misinterpreted as two independent biological signals without considering biosynthetic constraints. These intertwined dependencies, compositionality and biosynthetic constraints, mean that standard analytical approaches developed for gene expression data can produce misleading results when applied to glycomics datasets.

The standard solution in other omics fields, using simulated data with known ground truth to rigorously benchmark analytical methods^11–13^, has remained out of reach for glycomics. Existing simulation approaches^14,15^, typically relying on Dirichlet distributions, either generate unrealistic abundance patterns that fail to capture biological signal structure or cannot inject controlled experimental artifacts such as batch effects while maintaining compositional closure. Other parallel approaches to this model glycan biosynthesis^8,9,16^ and thus can achieve modeled abundance data, for either an improved understanding of biosynthesis or a tuning of the glycosylation of biologics, yet are unable to recapitulate the mentioned technical constraints of glycomics data.

One area of necessary method development in comparative glycomics data are batch effects^17^. Defined as the effect of external factors (e.g., sample storage duration, instrument differences, etc.) on the measurement, they manifest in changed inter-sample relationships, such as samples clustering by batch instead of by biological condition^18,19^. This results in both false-negatives and false-positives if these batch effects are not accounted for.

While normalization methods for glycomics data have been compared^20^, so far, the lack of readily available and realistic simulated glycomics data has prevented systematic evaluation of whether batch correction methods developed for transcriptomics^21^ translate to glycomics data, and, more fundamentally, whether the field’s growing arsenal of approaches for analyzing glycomics data can reliably distinguish biological signal from technical noise. Without the ability to generate realistic glycomics data, where the true differential signals and batch effects are known by design, researchers cannot objectively compare preprocessing pipelines, validate feature selection strategies, or establish evidence-based guidelines for when batch correction improves versus degrades analytical outcomes. Especially with the growing rise of efforts to identify diagnostic glycan-based biomarkers^5,22,23^, effective and robust data processing is key to prevent analytical errors.

Here we introduce GlycoForge, a simulation framework that operates in centered log-ratio (CLR) space to generate glycomics datasets with specified biological differences, controllable batch effects, and realistic mechanisms of simulating missing data, all while preserving compositional properties. We make GlycoForge available as a Python package as well as via a web interface at https://glycoforge.streamlit.app/ and provide example workflows as well as reproducible analyses of our findings at https://github.com/BojarLab/GlycoForge/blob/main/use_cases/glycoforge_analyses.ipynb.

GlycoForge supports two complementary modes: fully synthetic simulation from Gaussian copula sampling with covariance estimated from glycowork^24^ reference data for controlled hypothesis testing, and templated simulation that preserves the effect size geometry from real differential expression analyses while allowing for systematic variation of signal strength and technical artifacts. By injecting batch effects as directional shifts in CLR space before back-transforming to compositional abundances, GlycoForge maintains closure while enabling precise control over both the magnitude and structure of experimental biases. Both biological and batch effects can be specified on the motif level and use biosynthetic networks to create realistic shifts in glycomics data. We demonstrate GlycoForge’s utility by systematically evaluating batch correction methods such as ComBat^21^ and other methods, including newly developed variants, across a grid of biological signal strengths and batch effect severities, establishing quantitative criteria for when batch correction enhances versus obscures true biological differences in comparative glycomics studies. Beyond batch correction, GlycoForge provides a general-purpose testbed for any analytical workflow requiring ground-truth glycomics data in the form of processed abundance data, positioning the field to adopt the same rigorous, simulation-based method evaluation that has become standard practice in transcriptomics and related areas.

## Results

### GlycoForge generates compositionally valid glycomics datasets via CLR-space transformations

GlycoForge operates exclusively in centered log-ratio (CLR) space to maintain compositional closure while injecting controlled biological and technical effects. The CLR transformation maps compositional data from the simplex to an unrestricted Euclidean space where standard additive operations are valid^25^, enabling us to inject effects as simple arithmetic shifts before transforming back to valid compositional abundances via the inverse CLR. This mathematical framework ensures that all simulated datasets satisfy the fundamental constraint of relative abundances that glycan proportions sum to unity within each sample, preventing spurious correlations and invalid statistical inferences that arise from analyzing raw compositional data directly^5^. We note that the GlycoForge package exhibits basic plotting capabilities (such as the *plot_pca* function to obtain PCA biplots), yet figures shown in this manuscript have been in part obtained by functions/code found in Jupyter notebooks within the GlycoForge repository (https://github.com/BojarLab/GlycoForge) and were manually styled thereafter.

We implemented two complementary simulation modes to address different experimental needs (Fig. 1A). The synthetic mode generates fully simulated datasets from user-specified parameters without requiring input data, enabling controlled hypothesis testing with known ground truth, while the templated mode is guided by real input glycomics data. Starting from the baseline that Dirichlet sampling is routinely used in simulating glycomics data^14,15^, yet has never been systematically evaluated for this purpose, we first benchmarked several approaches in the templated simulation mode, by comparing distributional properties of simulated data to real glycomics data (Fig. S1). To our surprise, Dirichlet sampling was the worst tested alternative and a Gaussian copula of Ledoit-Wolf shrinkage covariance and empirical marginals from the used data template produced much more fitting simulated glycomics data, with better matching marginal distributions (median Kolmogorov-Smirnov statistic 0.29 vs. 0.53 for Dirichlet; Wilcoxon signed-rank p < 0.001) and correlation structures (median Mantel r > 0.75). For purely synthetic simulations, in the absence of a template, we automatically estimate these covariances from a pool of glycomics data of the specified glycan class, stored in the glycowork package.

**Figure 1.**
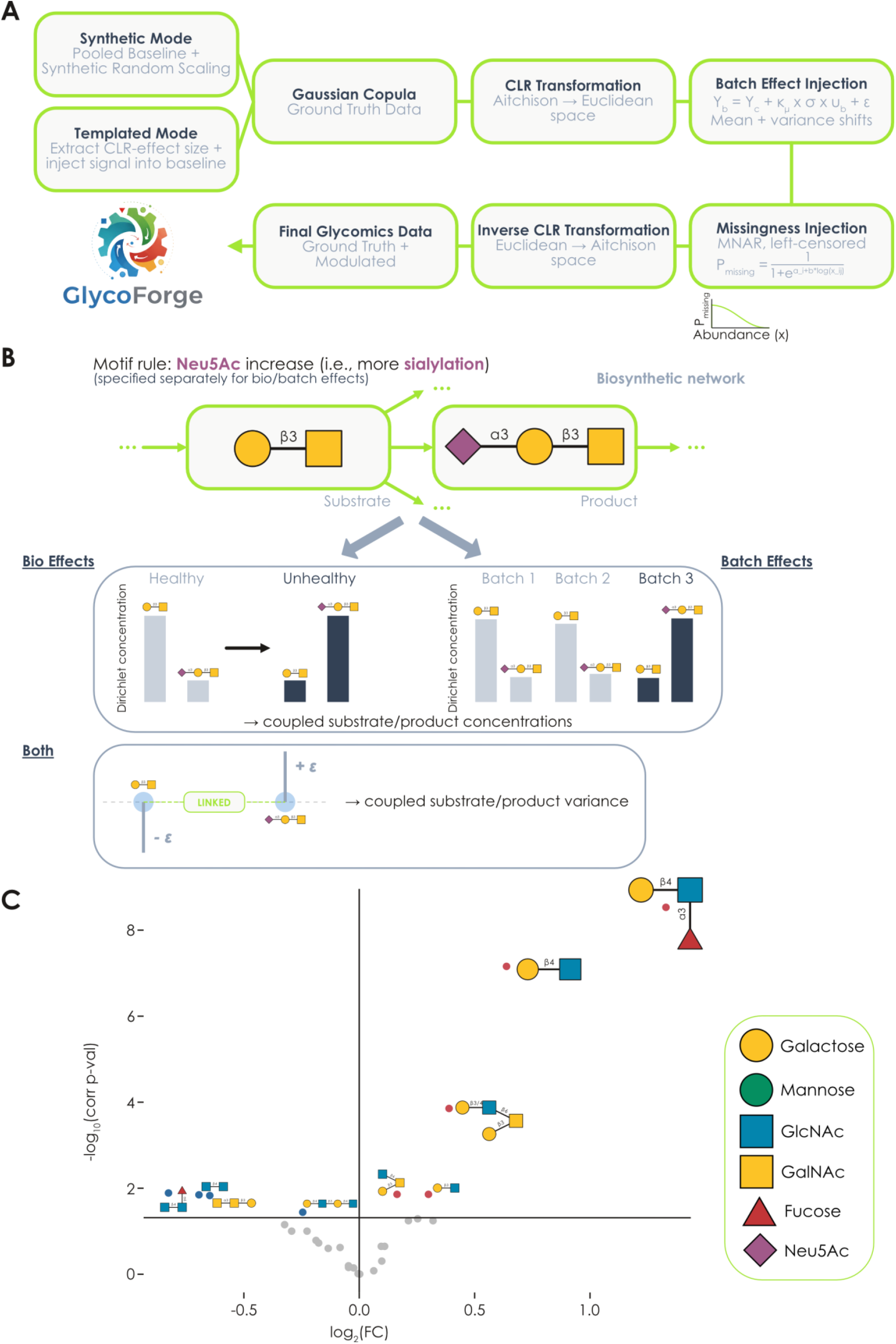
GlycoForge simulates fully synthetic as well as templated glycomics data with specified motif effects. **A)** Overview of the data simulation pipeline in GlycoForge, including both simulation modes as well as the functionality to, optionally, inject batch effects and/or realistic missing data. **B)** Injecting motif-level changes in differential abundances and batch effects. Based on user-provided motif-level changes for bio/batch effects, we retrieve relevant substrate/product relationships from a dynamically constructed biosynthetic glycome network^9^ and use those to redistribute proportions between substrates and products, yielding a CLR-space injection direction applied during Gaussian copula sampling. Further, substrates/products are variance-coupled to simulate their biosynthetic relationships. **C)** Introducing specific motif dysregulations into synthetic glycomics data. Shown is a motif-level volcano plot, generated with the *get_volcano* function from glycowork (v1.8.0), for synthetic data with 100 random *O*-glycans, drawn with the *get_random_glycan* function from glycowork, where Lewis A/X antigens, Fucα1-3/4(Galβ1-3/4)GlcNAc, were chosen to be upregulated. Glycans are drawn via GlycoDraw^27^ and conform to the Symbol Nomenclature For Glycans (SNFG).

For synthetic simulations, users specify the number of glycans and samples per group, and the pipeline constructs a healthy baseline proportion vector from the pooled mean abundance profile across qualifying glycowork reference datasets of the specified glycan class, then generates disease parameters through heterogeneous scaling where a specified fraction of glycans are randomly upregulated or downregulated by drawing scaling factors from configurable ranges. This stochastic heterogeneity produces realistic variability in effect sizes across features while maintaining complete control over the simulation parameters.

Glycome dysregulations are often changes on the motif level^5,14^, such as increased α2-3 linked Neu5Ac due to enzyme dysregulations, where up-/downregulations are spread across many glycans exhibiting that motif. To facilitate the simulation of such physiologically relevant effects, we expanded our simulation pipeline to also accept a dictionary of intended motif changes. Thus, users can provide specified effects on the motif level (e.g., increased sialylation), which are then used via a dynamically constructed biosynthetic network^9,26^ to redistribute proportions between substrates and products (Fig. 1B), to respect mass conservation. In this case, the products and substrates also share variance, to reflect their tight biological coupling. Concretely, for each substrate-product pair identified in the network, the substrate proportion is scaled down by a random factor and the lost mass is divided equally among its products, preserving the total composition exactly. The resulting perturbed proportion vector are then transformed into CLR space, and their difference δ = clr(pU) − clr(pH) serves as the biological injection direction that is added to the unhealthy rows during Gaussian copula sampling, so the motif-level flux redistribution governs the systematic CLR-space shift between groups.

The templated mode preserves authentic biological signal structure by starting from real glycomics data, extracting Cohen’s *d* effect sizes via differential expression analysis using the glycowork package (which has been developed for compositional glycomics data^5^), and injecting these effect size patterns into simulated data while allowing for systematic variation of signal strength and technical artifacts. This approach captures the complex correlation structure present in real biological differences while enabling rigorous method evaluation under known ground truth. Both simulation modes allowed for the simulation of large glycomics datasets in less than a second on a regular consumer laptop (Fig. S2A-B).

Relying on our benchmarking against real-world glycomics datasets, we next selected data normalization routines, and default values for these routines, that resulted in the best-matching glycomics data (Fig. S3). Raw Cohen’s *d* values extracted from differential expression analysis undergo a three-step processing pipeline that centers the distribution to remove global shifts, applies Winsorization to clip extreme outliers at automatically selected percentiles based on outlier severity, and normalizes by dividing by a baseline measure to produce standardized effect sizes suitable for scaling by the user-specified *bio*_strength_ parameter. These normalized effect sizes are then injected as additive shifts in CLR space with magnitude controlled by the *bio*_strength_ parameter, enabling systematic exploration of how biological signal strength interacts with technical artifacts such as batch effects. Effect size processing in the templated mode ensured robust biological signal injection while maintaining numerical stability (Fig. S4).

Synthetic data generated with 30% upregulation and 35% downregulation exhibit clear separation between healthy and unhealthy groups in principal component analysis (Fig. S5), with CLR-space injection vectors of randomly chosen glycan indices at varying magnitudes and directions, reflecting our implemented multi-eigenvector injection geometry that distributes biological signal broadly across the feature space. In contrast, templated data derived from a human GAG-transfected *N*-glycomics dataset^28^ show more structured effect size patterns that reflect coordinated biological processes (Fig. S5), where only altered glycans receive effect injections.

The CLR-space injection framework maintains several critical mathematical properties that ensure biologically valid simulations. First, compositional closure is guaranteed because all effects are injected in CLR space and converted back via inverse CLR, which by construction produces probability vectors summing to unity. Second, the zero-sum constraint of CLR in relative abundance data ensures that upregulation of some glycans necessarily implies relative decreases of others, which we actively leverage via our biosynthetic networks in synthetic mode. The Gaussian copula sampling framework derives inter-feature dependencies from the Ledoit-Wolf covariance of real reference data and preserves empirical marginal distributions via quantile mapping, producing realistic within-group heterogeneity without imposing parametric distributional assumptions. Finally, the separation of baseline distribution (copula correlation structure and empirical marginals) from effect injection (CLR-space shifts) allows for independent control of signal magnitude and biological variability, enabling systematic exploration of scenarios where strong biological effects coincide with either homogeneous or heterogeneous samples.

Since GlycoForge is built on top of the glycowork package^24^, this also allowed for powerful motif matching algorithms to be used for maximum flexibility in directing motif-level effects. As an example, we generated synthetic glycomics data with upregulated Lewis A/X antigens, Fucα1-3/4(Galβ1-3/4)GlcNAc, and show here that the target epitope was statistically significantly upregulated in the resulting glycomics data (Fig. 1C), along with epitopes co-occurring on Lewis A/X-containing glycans. Investigating whether motif injections also captured correlation structures of non-motif containing glycans (e.g., substrates being consumed to generate motif-containing products), we showed that templated (but not synthetic) simulations of Neu5Ac-dysregulation partly recovered such substrate-product correlations found in real glycomics data with this dysregulation (Fig. S6), albeit with a systematic underestimation of this effect strength (Pearson’s r 0.22 vs 0.47 in real data). Templated mode recovers these correlations because the CLR-space injection is derived from real differential expression effect sizes, which already encode the coordinated up- and downregulation of substrate-product pairs as it occurred in the original biological samples. Synthetic mode produces near-zero substrate-product correlations because its Gaussian copula is built from covariance pooled across all class-specific glycowork datasets by abundance rank rather than by glycan identity, so the covariance at any pair of feature positions reflects the average correlation between abundance-ranked slots, not the biosynthetic relationship between the specific glycans assigned to those positions. When possible, we therefore recommend users to use templated simulations.

### GlycoForge can inject data imperfections such as batch effects or missingness into compositional glycomics data

Realistic evaluation of analytical methods requires simulated data that recapitulates the technical artifacts present in real experimental measurements. GlycoForge addresses this by implementing two major sources of data imperfection commonly encountered in glycomics studies: batch effects arising from sources such as sample processing heterogeneity and missing values reflecting detection limit constraints in mass spectrometry measurements. Both artifact types are injected in ways that maintain compositional closure while producing statistically detectable and biologically plausible distortions of the underlying signal.

Batch effects in GlycoForge are implemented as sparse directional shifts in CLR space, where each batch receives a unique direction vector specifying which glycans experience systematic displacement and in which direction. Here again, we provide the option to direct the type of batch effect per batch on the motif-level, such as decreased sialylation in some batches (Fig. 1B). This simulates realistic processes, such as sialylated glycans decreasing in abundance over extended periods of storage^29,30^.

The direction vectors then are normalized to unit magnitude to ensure consistent batch effect strength independent of the number of affected glycans, and the same direction structure is maintained across all parameter combinations within a simulation study to isolate the effects of varying batch severity from confounding changes in batch geometry. The actual batch effect applied to each sample combines two components: a mean shift term that systematically displaces samples along the batch direction proportional to the baseline glycan-wise standard deviation and controlled by the *κ*_*μ*_ parameter, and a variance inflation term that changes variance between batches, scaled by a batch-specific multiplier controlled by the *var*_*b*_ parameter. This formulation creates both location and scale heterogeneity across batches, while maintaining proportionality to the natural variability of each glycan, producing batch effects that scale appropriately with feature abundance patterns rather than imposing uniform absolute shifts that would be unrealistic for compositional data.

The impact of batch effects on data structure becomes visually apparent in principal component analyses as batch severity increases (Fig. 2A). At low batch effect strength, samples retain clear clustering by biological status with minimal batch-induced displacement, and the biological effect size measured by η^2^ remains dominant at 0.72. As batch parameters increase, however, batch-specific clusters become visible within each biological group. At the highest severity tested, batch completely dominates the variance structure with a η^2^ of 0.84 versus only 0.05 for biological status, producing PCA plots where samples cluster primarily by batch rather than by disease state. This progression demonstrates that GlycoForge can systematically span the full range from negligible technical noise to severe confounding where batch effects overwhelm biological signal, enabling rigorous evaluation of correction methods across the artifact severity spectrum encountered in real glycomics studies.

**Figure 2.**
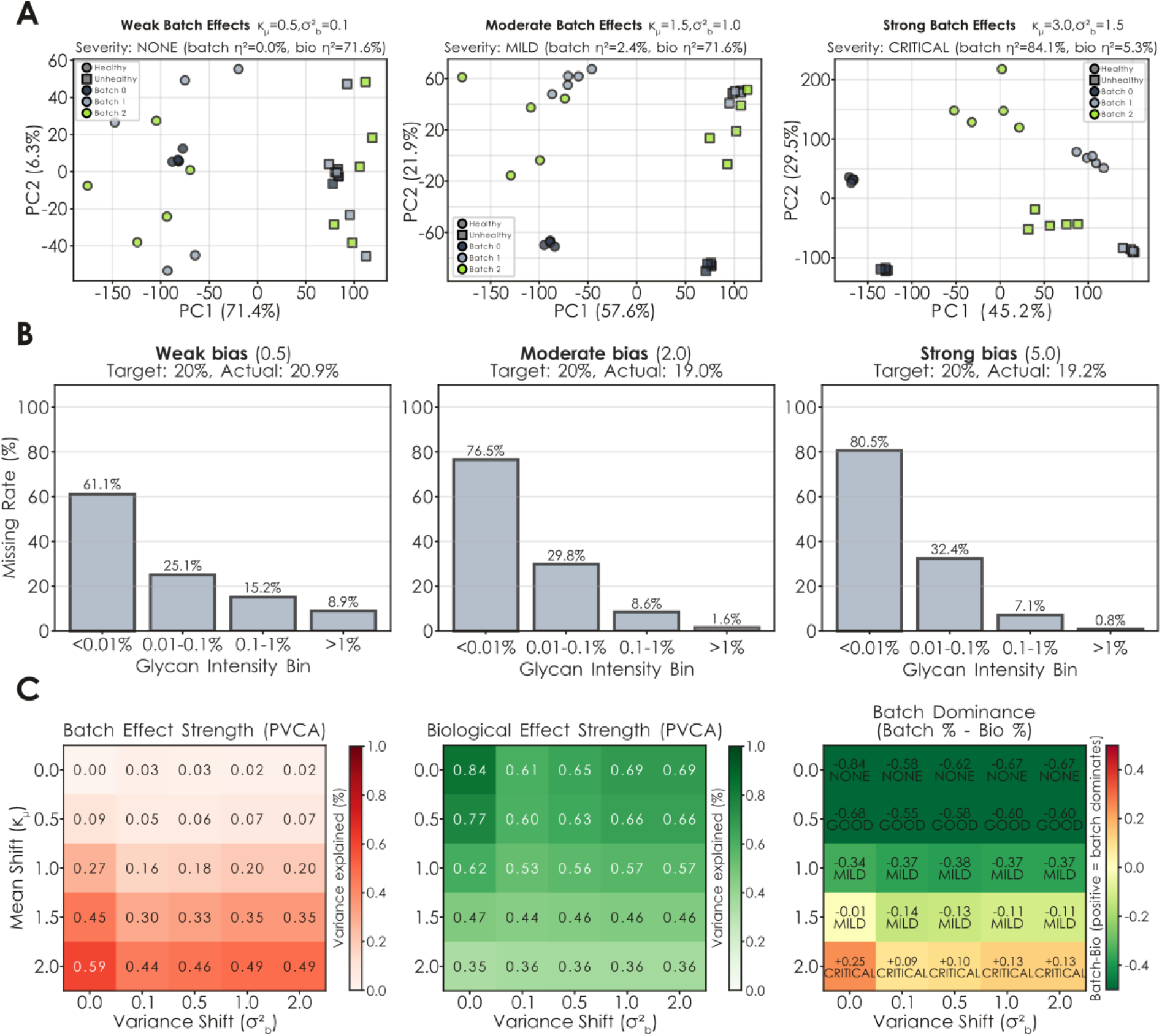
Glycomics data with desired imperfections can be simulated with GlycoForge. **A)** Visualizing batch effects introduced into synthetic glycomics data. Shown are PCA plots of simulated glycomics data (n = 15 per condition) with varying batch effect strengths. Biological status is indicated by shape, whereas batch is indicated by color. Mean and variance shifts are provided, as well as the effect size of batch and biological status as η^2^. **B)** Evaluating left-censored bias in introducing missing data in synthetic glycomics data. Bar graphs of the distribution of chosen missing values across intensity bins are shown. Stronger biases favoring left-censored missingness resulted in a higher proportion of missing values being drawn from less abundant glycans. **C)** Assessing batch effects in synthetic glycomics data, with increasing mean and variance shifts. Batch and biological effect were assessed via their respective percent-variance-explained of a principal variance component analysis (PVCA). Batch dominance was then quantified as the difference between these two (Batch % - Bio %).

Missing data in glycomics arise predominantly from abundance-dependent detection limits in data-driven acquisition (DDA) mass spectrometry^31^, where low-abundance glycans are more likely to fall below quantification thresholds because of detection limits. GlycoForge implements this left-censored missing-not-at-random (MNAR) pattern by computing sample-specific missingness probabilities as a function of normalized glycan intensity. We analyzed real-world glycomics data to determine a logistic decrease, with a slope of b = 1.0, as the best pattern on average to simulate realistic missingness that generalized across glycan classes and laboratories (Fig. S7; Wilcoxon signed-rank p < 0.001, n = 24 glycomics datasets). Users can also modulate this slope parameter within GlycoForge to control the strength of the low-abundance bias. Within each sample independently, glycans are assigned missing status according to Bernoulli draws using these intensity-dependent probabilities, where a per-sample intercept is solved numerically so that the expected missing fraction matches the user-specified target within each sample. This approach ensures within-sample missing patterns that matches real-world glycomics data properties.

The effectiveness of the intensity-dependent missingness model is evident in the distribution of missing values across abundance bins (Fig. 2B). Even under weak bias (b = 0.5), the missing rate decreases monotonically with glycan intensity, from 61.1% in the lowest abundance bin to 8.9% in the highest, confirming that the logistic model produces left-censored patterns at all parameter values. As bias strength increases to the empirically calibrated default 1.0 and beyond, the left-censored pattern becomes increasingly concentrated in the lowest abundance bins, with progressively fewer missing values in higher abundance ranges, closely mimicking the systematic loss of low-intensity glycans observed in real mass spectrometry data (Fig. S7A-B). This tunable intensity-dependence ensures that imputation strategies are evaluated under realistic data loss scenarios. We note that both missing data and batch effect injection hardly increase runtime of simulations (Fig. S1C).

The interplay between biological signal strength and severity of imperfections such as batch effects can also be quantified comprehensively using principal variance component analysis (PVCA)^32^, which decomposes variance across multiple principal components rather than focusing on a single axis (Fig. 2C). At low batch severity, biological variance dominates across all tested biological signal strengths, with batch variance barely exceeding 10%. Notably, the mean shift parameter (*κ*_μ_) drives batch severity almost entirely, while increasing the variance shift (*σ*^2^_b_) does not strengthen the batch effect but instead adds isotropic within-batch noise that dilutes both batch and biological signal. As the mean shift increases, a critical transition occurs where batch variance begins to exceed biological variance, especially when biological effects are weak. This batch dominance metric proves particularly useful for establishing evidence-based guidelines about when batch correction is necessary, as correction methods should primarily be applied when batch effects detectably distort the variance structure rather than being used indiscriminately regardless of batch severity, risking overcorrection. We thus provide this functionality as the *glycoforge*.*utils*.*check_batch_effect* function to users during data exploration if they have different sets of samples (e.g., stored for different amounts of time, or measured on different instruments).

### Benchmarking batch effect correction methods for simulated and real-world glycomics data

Next, we used the GlycoForge platform to systematically benchmark different batch effect correction methods, including two variants newly developed here, to determine the current state-of-the-art for compositional glycomics data. We evaluated six batch correction methods across a parameter grid spanning biological signal strengths (*bio*_strength_ from 0.5 to 2.0) and batch effect severities (*κ*_*μ*_ from 0.0 to 2.0, *var*_*b*_ from 0.0 to 1.0), including a baseline condition without batch effects, using synthetic glycomics data with three random seeds per condition. The tested methods represent diverse algorithmic strategies: ComBat^21^ applies empirical Bayes shrinkage to location and scale parameters while protecting biological covariates, percentile normalization maps feature distributions to a reference batch, ratio-preserving ComBat (developed here) modifies standard ComBat to operate in log-ratio space with explicit renormalization, Harmony^33^ uses iterative clustering in PCA space, limma-style correction^34^ applies linear regression to estimate and subtract batch effects, and stratified ComBat (developed here) applies standard ComBat independently within each biological group.

Overall, standard ComBat and Ratio-ComBat emerged as the clearly superior methods in terms of removing the batch effect (Fig. 3A). Both methods reduced PVCA batch variance to near zero, while other methods such as Percentile or Harmony barely reduced the batch effect. Similar patterns held across other batch effect metrics.

**Figure 3.**
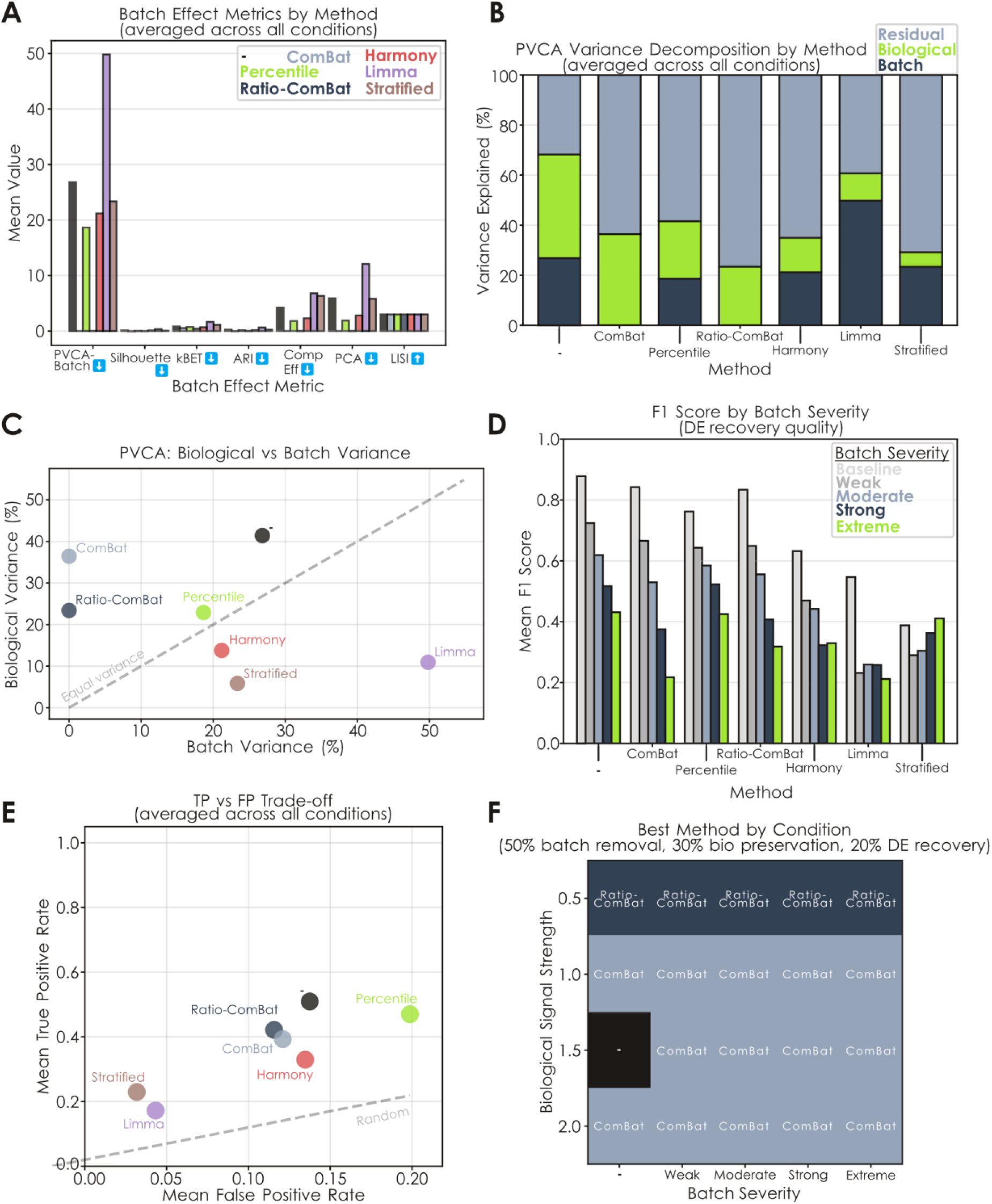
ComBat batch effect correction is effective for glycomics data. **A)** Assessing different batch effect correction methods on synthetic glycomics data. Across a grid of biological effect and batch effect strengths (both mean and variance shifts), and repeated with three different seeds, a range of metrics were evaluated for each method (averaged across all conditions) and are shown via bar graphs. The arrow indicates whether a metric is supposed to be low or high. **B)** Explained variance by condition via a principal variance component analysis (PVCA) after batch effect correction. For each method, the percent-variance-explained of biological status, batch effect, and residual variance is shown as a stacked bar graph. **C)** Comparing explained biological variance and batch variance for the different metrics. A trend line for the two being equal is shown in grey. Methods above the line exhibit higher biological signal than batch signal. **D)** Comparing batch effect correction methods to their ability of retaining significant differences prior to batch effect introduction. Averaged F1 scores for each batch effect severity level and each method are shown as bar graphs. **E)** Comparing true-positive and false-positive rates after batch effect correction. A trend line for equal true- and false-positive rates is shown in grey. Methods above the line exhibit better-than-random performance. **F)** Aggregate metric to choose batch effect correction method by batch severity. For every combination of biological signal strength and batch severity, the best correction method was chosen by weighting their impact on batch effect removal and biological signal preservation.

Removing a batch effect is only worthwhile if overcorrection does not remove biological signal. PVCA revealed that stratified ComBat removed nearly all biological signal while leaving substantial batch variance intact (Fig. 3B-C). This indicates that global correction with biological covariate protection outperforms alternatives such as the within-group correction strategy. The other methods largely preserved biological effects, with ComBat, Percentile, and Ratio-ComBat exhibiting the best performance. However, for Percentile, Harmony, limma, and stratified ComBat, batch variance continued to exceed biological variance even after correction, suggesting undercorrection. We note that methods such as limma and Harmony have been developed for transcriptomics data and, in the case of Harmony, even single-cell transcriptomics data. Limma is further primarily a differential expression platform, rather than a batch effect removal pipeline.

Differential expression recovery through increasing batch effect strengths, quantified via F1 scores, also demonstrates that stratified ComBat severely degrades statistical power (Fig. 3D). Here, Percentile, Harmony, and limma exhibited the most robust response to extreme batch effects, while ComBat and Ratio-Combat were sensitive up to large batch effects but overcorrected in the case of extreme batch effects. On average, Percentile normalization exhibited the highest false-positive rate among all methods, while limma and Stratified ComBat had the lowest false-positive rates but also the lowest sensitivity (Fig. 3E). We further note here that methods such as limma do not account for compositional data characteristics and in general are known to introduce high false-positive rates when applied to such data^35,36^. ComBat and Ratio-ComBat achieved the best trade-off between true-positive and false-positive rates among the tested methods. It should be noted that, on average, all methods were more likely to introduce false-positives under weak batch effects, while false-negatives were enriched in strong batch effects (Fig. S8), demonstrating the potential for overcorrection in either direction and demanding a guide for when to use a batch effect correction approach, which we provide via the *glycoforge*.*utils*.*check_batch_effect* function that classifies the impact of the batch effect via PVCA as shown in Fig. 2C, if suspected batch labels are known.

For a final method benchmarking, we integrated multiple evaluation dimensions into a composite decision framework to evaluate (i) batch effect removal as well as (ii) biological effect preservation. We performed a grid search of different weightings of these aspects (all weight combinations in 10% steps). An example outcome, weighted 50% batch variance removal, 30% biological variance preservation, and 20% differential expression recovery, can be seen in Fig. 3F. Across nearly all combinations of biological signal strength and batch severity, standard ComBat emerged as the optimal method in 63.9% of weight combinations, followed by limma (16.8%) and Ratio-ComBat (10.8%). Ratio-ComBat became preferred only with weak biological effects (Fig. 3F), where it provided a marginal advantage over standard ComBat. No other method consistently won under any condition, predominantly due to insufficient batch effect removal. However, we note that this superior performance of ComBat may be specific to additive batch effects, as we identified a preference for uncorrected data in the case of multiplicative batch effects (i.e., non-linear effects where the batch effect strength is modulated by signal intensity; Fig. S9).

We next confirmed ComBat’s performance on a more domain-specific batch effect and simulated the loss of sialylation in some batches (Fig. S10) as well as injected batch effects into real-world glycomics data (Fig. S11). Lastly, we could also show that inter-lab differences for glycomics data of the same tissue and glycan class can be modeled as a batch effect and efficiently removed via ComBat without losing biological signal (Fig. S12). In all cases, ComBat was still able to remove the batch effect, confirming its suitability for glycomics data. We however caution that high degrees of confounding between batch and bio group memberships make batch effect removal unfeasible (Fig. S13). The superior performance of standard and ratio-preserving ComBat validates the CLR-space correction paradigm and demonstrates that methods originally developed for gene expression microarrays translate effectively to compositional glycomics data when applied with appropriate covariate protection.

## Discussion

The difficulty of experimentally collecting large sets of glycomics data with specified group differences, coupled with a plethora of downstream analysis methods, means that there is a need for simulating realistic glycomics data and its imperfections. We here present such a platform with GlycoForge, which provides users with full control over experimental imperfections (e.g., batch effects, missing data) as well as group differences (including realistic motif-level dysregulation). All this is (i) fully seeded for reproducibility, (ii) inherently compatible with the compositional nature of glycomics data, and (iii) performant enough for large-scale simulations on regular laptops. We adopt a Gaussian copula as the simulation backbone of GlycoForge based on its empirically validated superiority over Dirichlet sampling for glycomics data, though future approaches may improve upon this further.

GlycoForge is modular and functionalities such as the missing data injection can be used or not in any simulation workflow. We are confident that the open-source nature of our package will lead to further extensions by us and the community. We also see great potential in such future developments of GlycoForge’s capabilities, beyond the scope of this work. One avenue here would be an extension of the templated simulation to settings beyond two-group comparisons, given emerging glycomics datasets of multiple cancer stages^37^ or infections with various pathogens^38^. Currently, there is also a lack of temporal analysis approaches for glycomics data that have been rigorously assessed, while data of this type are being generated^39^, which presents another opportunity. Further, we note that GlycoForge is designed to produce processed glycan abundances and future work could extend it to the simulation of the original glycan ion intensities as well, e.g., to facilitate benchmarking of data processing workflows. Finally, future research could also explore ways to use the mechanistic control provided by our biosynthetic modeling to use, e.g., glyco-transcriptomic data for biologically relevant perturbations and simulated data that could then be used for both benchmarking and even hypothesis generation. This could then also extend our usage of biosynthetic networks and shifting motif-level flux, which we currently restrict to direct substrate-product relationships that could be generalized by future work.

Regarding our experiments on batch effect correction for glycomics data, we report that ComBat is, currently, the best overall approach, with Ratio-ComBat being applicable in the case of weak biological effects. However, we caution that (i) ComBat introduces false-positives in the case of weak batch effects and (ii) sensitivity is partially lost with ComBat with strong batch effects. We thus argue that future work could use our benchmarking platform to develop more suitable algorithms for glycomics data to improve batch effect correction further, especially given the poor performance ComBat exhibited with multiplicative batch effects. Further, it should be noted that batch effect correction is only possible if batches contain samples from all conditions, as otherwise the batch effect is inextricably confounded with biological effects. In addition to this, researchers have to know the batch variables and which samples belong to which batch, to apply methods such as ComBat, whereas surrogate variable analysis^40^ is required for cases without this knowledge.

Overall, we view GlycoForge as a one-size-fits-all solution to simulating glycomics data that can be further expanded, aided by our open-access approach, as the needs of the community shift. Built on top of the glycowork platform, GlycoForge has access to advanced motif annotation^41^, glycan nomenclature flexibility^42^, and differential expression capabilities^5,14^, which also will grow in performance over time. Next to bioinformatics-related needs described throughout this work, we expect end users (aided by our web app https://glycoforge.streamlit.app/) to use GlycoForge for applications such as estimating batch effect strengths in collected data or during study design when determining sample size for the reliable detection of expected motif-level effects (Fig. S14). We envision that GlycoForge will be used to establish a robust foundation for comparative glycomics workflows of the future, such as we have already shown here with the problem of batch effect correction, yielding reliable biomarker candidates and understanding glycome dysregulation in disease.

## Methods

### GlycoForge simulation framework

GlycoForge (v0.2.2) is a Python-based simulation pipeline for generating glycomics relative abundance datasets with controllable biological group differences and injectable batch effects as well as data missingness. The framework operates exclusively in centered log-ratio (CLR) space to maintain compositional closure throughout all transformations. All code is available at https://github.com/BojarLab/GlycoForge and requires Python ≥3.10 with core dependencies glycowork≥1.8.0, NumPy, pandas, SciPy, scikit-learn, Matplotlib, and seaborn.

### Centered log-ratio transformations and compositional data handling

Glycan relative abundances are compositional data constrained to sum to 100%, residing on the simplex. Direct manipulation of compositional data violates independence assumptions and induces spurious correlations^5^. To address this, GlycoForge implements the centered log-ratio transformation for all effect injections. For a compositional vector **x** with *n* features, the CLR transformation is defined as

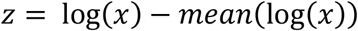

where ‘mean’ refers to the geometric mean, which ensures the transformed data lie in a (*n*-1)- dimensional Euclidean space where standard additive operations are valid. The inverse CLR transformation is computed as

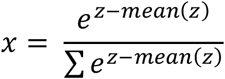

with an optional scaling to percentages by multiplying by 100. For numerical stability, the CLR vector is centered and then shifted by its maximum before exponentiation and zero values are replaced with 10^−6^ before log transformation. The CLR framework ensures that all simulated batch effects and biological signals can be injected as additive perturbations in z-space without violating the simplex constraint, and the inverse CLR guarantees a return to valid compositional data.

### Synthetic simulation mode

The synthetic simulation mode generates glycomics datasets without requiring data input from the user. Inter-glycan correlation structure is learned from real glycowork reference data and injects user-defined perturbations. The user specifies the number of glycans *n* and sample sizes *N*_H_ (healthy) and *N*_U_ (unhealthy). The pipeline loads all glycomics datasets from glycowork matching the specified glycan class (*N, O*, or GSL) and containing at least *n* glycans. For each qualifying dataset, the top *n* glycans by mean abundance are selected, samples containing zeros are removed, CLR transformation is applied, and a Ledoit-Wolf shrinkage covariance matrix is estimated. Per-dataset correlation matrices are averaged to form a consensus correlation matrix R, and marginal standard deviations are pooled across all datasets to reconstruct a covariance matrix

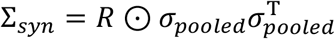

where ⊙ denotes element-wise multiplication and the outer product σ_pooled_ σ_pooled_^T^ gives the matrix of pairwise standard deviation products. The healthy baseline proportion vector *p*_H_ is constructed from the pooled mean abundance profile across qualifying datasets, normalized to sum to 1, and used to define the healthy weighted concentration vector *α*_H_ = 10*n*\**p*_H_. For each random seed, the unhealthy concentration *α*_U_ is generated through heterogeneous scaling of *α*_H_. By default, 30% of glycan indices are randomly selected for upregulation without replacement, and each receives a scaling factor uniformly sampled from [1.1, 3.0]. Then, 35% of the indices are selected for downregulation, each receiving a scaling factor from [0.3, 0.9]. The remaining 35% of glycans are unchanged between groups. The scaling is applied as

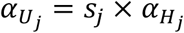

for affected glycans, where *s*_*j*_ is the respective scaling factor, and *α*_U_ is clipped to a minimum of 10^−3^ to ensure numerical stability. Clean compositional data are then generated by drawing Gaussian samples Z ∼ 𝒩(0, R_syn_ + εI), where ε = max(10^−8^, 10^−4^ · *n*), applying the probability integral transform U = Φ(Z) to obtain uniform marginals, and mapping these uniforms to empirical CLR marginals via linear interpolation through the sorted pooled synthetic CLR reference matrix. When motif rules are provided, the biological injection direction is computed as δ = clr(pU) − clr(pH) and added directly to the *N*_U_ unhealthy rows When no motif rules are provided, the injection is constructed from the top-K eigenvectors (K = min(3, n-1)) of Σ_syn_ with independent random signs, and each eigenvector direction 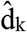 is scaled as 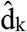· λ · s_k_ · √n, where λ is the *bio*_strength_ parameter and s_k_ is the standard deviation of the CLR data projected onto 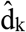. This multi-eigenvector injection ensures the biological signal is distributed across multiple principal components and remains detectable via PVCA at *bio*_strength_ values that are interpretable as standard effect sizes. The injected CLR data are then transformed back to percent-scale compositions via softmax.

### Templated simulation mode with real data effect integration

The templated mode preserves authentic biological signal structure by starting from real glycomics data. The user provides a CSV file containing relative abundance measurements and specifies column prefixes to identify healthy and unhealthy sample groups. Differential expression is computed using glycowork’s *motif*.*analysis*.*get_differential_expression* function with CLR transformation and imputation enabled^5,14,24^, yielding Cohen’s *d* effect sizes for each glycan. Because glycowork may filter out some glycans during analysis, the returned effect sizes are reindexed to match the original glycan order, with missing glycans assigned effect sizes of zero to indicate no injected effects. The differential mask parameter controls which glycans receive biological signal injection and accepts either string specifications (“All”, “significant”, “Top-N”) or custom binary arrays. The “significant” mode uses only glycans marked as statistically significant by glycowork’s sample-size-adjusted alpha and multiple testing correction. The “Top-N” mode selects the N glycans with largest absolute effect sizes. The healthy baseline composition *p*_H_ is extracted as the column mean across all healthy samples, and any zero values are imputed as 10% of the minimum nonzero value (or 10^−3^ if all values are zero) before normalizing to sum to 1. This baseline is transformed into CLR space as *z*_H_ = clr(*p*_H_). The effect sizes undergo robust processing through centering via

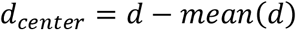

followed by Winsorization where outliers are clipped to a percentile threshold that is automatically selected based on outlier severity (measured as max absolute value divided by 75^th^ percentile). If this ratio exceeds 15, the 85^th^ percentile is used; if it exceeds 10, the 90^th^ percentile; if it exceeds 5, the 95^th^ percentile; otherwise the 99^th^ percentile. After clipping, a baseline is calculated using either the median of absolute values (default), the median absolute deviation (MAD) scaled by 1.4826, or the 75^th^ percentile of absolute values. The processed effect sizes *d*_robust_ (obtained by dividing the Winsorized effects by this baseline) are then injected in CLR space as

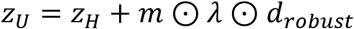

where **m** is the differential mask binary vector, *λ* is the *bio*_strength_ parameter controlling signal intensity, and ⊙ denotes element-wise multiplication. The unhealthy baseline proportions are recovered via inverse CLR as *p*_U_ = invclr(*z*_U_). Concentration parameters are constructed with variance control as

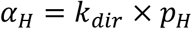

and

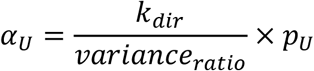

where *k*_dir_ controls overall concentration (inversely related to variance) and *variance*_ratio_ ≥ 1 allows the unhealthy group to exhibit greater heterogeneity than the healthy group. Both *α*_H_ and *α*_U_ are clipped to [0.5, *α*_max_] to ensure valid concentration parameters. Clean compositional data are then generated using the same Gaussian copula procedure described for synthetic mode, substituting a Ledoit-Wolf shrinkage covariance estimated from the pooled CLR-transformed real samples (both healthy and unhealthy) for Σ_syn_, and using the real CLR data as the empirical marginal reference. The biological injection vector δ = m ⊙ λ · *d*_robust_, where **m** is the differential mask and *d*_robust_ the processed effect sizes, is added to the *N*_U_ unhealthy rows before the final softmax transformation.

### Batch effect direction vector generation

Batch effects in GlycoForge are defined as sparse direction vectors in CLR space. The user can provide fixed batch-glycan mappings or allow random generation controlled by parameters. In random mode, for each of *N*_b_ batches, the number of affected glycans is sampled from a normal distribution with mean (*f*_min_ + *f*_max_)/2 and standard deviation (*f*_max_ − *f*_min_)/4, clipped to [*f*_min_, *f*_max_] and converted to an integer count, where *f* is the affected fraction. The default range of *f* is [0.05, 1.00], meaning each batch affects 5-100% of glycans. With overlap probability *p*_o_ (default 0.5), glycans are selected from the full set allowing overlap between batches; otherwise, they are selected from glycans not yet affected by previous batches to minimize overlap. For each affected glycan in batch b, a direction *d*_bj_ is assigned as +1 with probability “positive_prob” (default 0.6) or -1 otherwise. These directions are assembled into a sparse vector *w*_b_ with *w*_b[j−1]_ = *d*_bj_ for affected glycans and 0 elsewhere. The vector is normalized by first centering to zero mean as

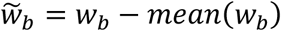

and then dividing by the root mean square norm to produce the final unit-normalized direction vector

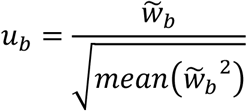

This normalization ensures batch effects are compositionally neutral in expectation and have standardized magnitude regardless of the number of affected glycans. The random generation uses a fixed seed (default 42) to ensure reproducible batch effect structure across different simulation runs, isolating variability to biological sampling and batch assignment rather than the batch effect definition itself.

### Stratified batch assignment and effect injection

Samples are assigned to batches using stratified random sampling to maintain biological balance within each batch. For each biological group (healthy and unhealthy), sample indices are shuffled and split into *N*_b_ approximately equal parts. Batch labels are assigned such that each batch contains a proportional mix of healthy and unhealthy samples. This stratification prevents complete confounding of batch with biology, reflecting realistic experimental designs where technical batches process multiple biological conditions. The baseline variance σ for each glycan is estimated from the clean CLR-transformed data as the standard deviation across all samples. For each sample *i* assigned to batch *b*_*i*_, the batch effect is applied in CLR space to the clean data *Y*_c_ as

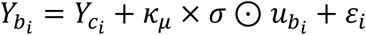

where *κ*_μ_ is the mean shift strength parameter, *σ* is the vector of per-glycan baseline standard deviations, *u*_*bi*_ is the batch direction vector for batch *b*_*i*_, and *ε*_*i*_ is sampled from a multivariate normal distribution

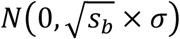

where *s*_*b*_ is a batch-specific variance scale factor: for *n*_*b*_ batches, batch *b* receives scale *s*_*b*_ = max(0.1, 1.0 + offset_b_) where

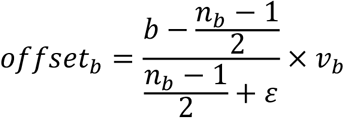

distributing batch-specific variance multipliers evenly around unity. This formulation creates moderate heteroscedasticity across batches while scaling batch effects proportionally to the natural variability of each glycan and maintaining independence across glycans. The mean shift term creates systematic displacement along the batch direction, while the variance term adds random noise that inflates spread within batches. After batch effect injection, the data remain in CLR space for downstream analysis or are transformed back to compositional space via inverse CLR for evaluation.

In addition to the default additive mode, GlycoForge supports a multiplicative batch effect mode where the mean shift is proportional to the current CLR value of each sample rather than being a constant displacement. In multiplicative mode, the mean shift for sample *i* is computed as (exp(*κ*_μ_ · *u*_*b*_) − 1) · *Y*_*ci*_, making the batch perturbation signal-dependent and producing non-linear distortions. The variance inflation term in multiplicative mode similarly scales with the absolute CLR value rather than the baseline σ, specifically using max(|*Y*_*ci*_|, 0.1·σ) as the noise scale.

### Missing data simulation

Broadly inspired by prior work on proteomics data^13^, missing data were simulated using a left-censored, missing-not-at-random (MNAR) pattern that reflects the intensity-dependent detection limits observed in real glycomics data. For each sample *i* independently, the missing probability for glycan *j* follows a logistic model in log-abundance space, where x_ij_ denotes the abundance of glycan *j* in sample *i, b* is the *mnar_bias* parameter controlling the steepness of the intensity-detection relationship (default *b* = 1.0), and *a*_i_ is a per-sample intercept calibrated individually for each sample, calculated as

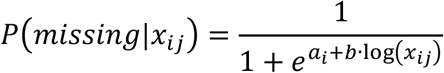

The intercept *a*_i_ is solved numerically using Brent’s method on the interval [-50, 50] to enforce the constraint that the mean missing probability across all glycans in sample *i* equals the target missingness fraction as best as possible. Binary missingness indicators were then sampled from Bernoulli distributions with the resulting per-glycan probabilities. Missingness was applied after batch effect injection to reflect the data acquisition process where missing values arise during measurement. Missing values were represented as NaN in compositional space and imputed with a detection limit value (1×10^−6^) prior to centered log-ratio transformation to enable downstream statistical analysis.

### ComBat batch correction implementation

Batch correction is applied using the ComBat algorithm^21^ implemented in Python, following the empirical Bayes framework originally developed for microarray data. ComBat takes as input the CLR-transformed data matrix (features × samples), batch labels, and an optional design matrix encoding biological covariates to protect. For this study, the design matrix is constructed from biological group labels using one-hot encoding to ensure ComBat preserves known healthy versus unhealthy differences during batch effect removal. The algorithm first fits a linear model to estimate overall biological effects as

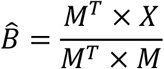

where *M* is the design matrix and *X* is the data matrix. The grand mean is computed as

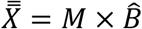

while residuals

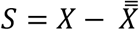

are further standardized by feature-wise standard deviations (*sds*), resulting in *S*^*^. For each batch *b* and feature *j*, location parameters 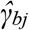 (mean batch effect) are estimated as the mean of standardized residuals in batch *b*, and scale parameters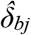 (variance batch effect) are estimated as the variance of residuals around the batch-specific mean. In parametric mode (used in this study), empirical Bayes shrinkage is applied to stabilize estimates for small batches. Prior hyperparameters are estimated from the distribution of batch effect estimates across batches, and posterior estimates are computed by combining the observed batch-specific estimates with these priors. The shrinkage is particularly important for small batches where empirical estimates are noisy. Finally, batch-corrected data are obtained by

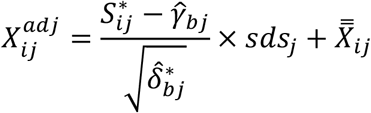

where 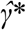 and 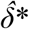 are the shrunken estimates and *S*\* is the standardized residual matrix. This transformation removes batch-specific mean shifts and variance inflations while restoring the original feature-wise scales and preserving the biological signals encoded in the design matrix.

### Other evaluated batch effect correction methods

In addition to ComBat, we evaluate five alternative batch correction methods to assess performance across different algorithmic strategies. Percentile normalization preserves compositional structure by normalizing feature distributions to a reference batch without requiring CLR transformation. For each sample, features are ranked and their values mapped to corresponding percentiles in the reference batch distribution, maintaining within-sample rank relationships while harmonizing cross-batch distributions. Ratio-preserving ComBat is a novel modification of the standard ComBat framework designed to respect compositional constraints by operating in log-ratio space with mean-centering at each step and transforming corrected values back to the simplex via exponentiation and renormalization, ensuring the sum-to-one constraint is maintained throughout. Harmony^33^ uses iterative clustering in PCA-reduced space, computing batch-specific cluster centroids and progressively shifting each batch toward global centroids while preserving local neighborhood structure. Limma-style correction^34^ applies a simpler linear model approach where batch effects are estimated as regression coefficients from a design matrix containing batch indicators, and batch-corrected data are obtained by subtracting the estimated batch effects from the original data followed by renormalization. Stratified ComBat is a variant that applies standard ComBat independently within each biological group (healthy versus unhealthy), preventing the algorithm from inadvertently removing biological differences when correcting for technical variation, which is particularly relevant when batch effects and biological effects are confounded or when biological groups exhibit different compositional baselines.

### Batch effect quantification metrics

Multiple complementary metrics quantify batch effect severity before and after correction. Principal Variance Component Analysis (PVCA) provides comprehensive variance decomposition by analyzing the first N principal components (default N = 10). For each *PC*_*i*_ with explained variance weight *w*_*i*_, the variance is partitioned into batch, biological, and residual components via ANOVA. The batch variance contribution is calculated as

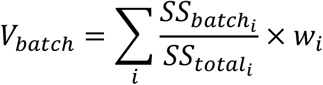

where *SS*_*batch,i*_ represents the sum of squares attributed to batch effects in *PC*_*i*_. Similarly, *V*_bio_ and *V*_residual_ are computed for biological and residual variance. The final percentages are normalized as

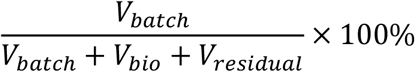

PVCA serves as the primary metric for overall batch effect severity assessment, as it captures variance contributions across multiple PCs.

The silhouette score measures cluster separation based on batch labels, computed as

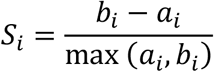

where *a*_*i*_ is the mean distance from sample *i* to other samples in the same batch and *b*_*i*_ is the mean distance to samples in the nearest different batch. Scores range from -1 to +1, with higher values indicating stronger batch separation and values near 0 indicating no batch structure.

The k-nearest neighbor batch effect test (kBET) evaluates whether the k-nearest neighbors of each sample have the expected batch composition. For each sample, a chi-squared statistic

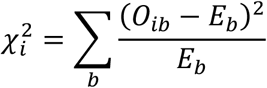

is computed, where *O*_*ib*_ is the observed count of batch *b* neighbors and *E*_*b*_ is the expected count based on global batch proportions. The mean chi-squared statistic across all samples serves as the kBET score, with higher values indicating departure from random batch mixing.

The Local Inverse Simpson’s Index (LISI) measures batch diversity in local neighborhoods by computing

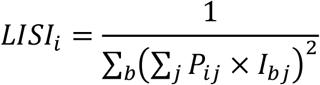

where *P*_*ij*_ is a kernel-weighted probability based on Euclidean distance and *I*_*bj*_ indicates batch membership. LISI values closer to the total number of batches indicate better mixing.

The Adjusted Rand Index (ARI) compares k-means clustering results to batch labels, measuring agreement between discovered clusters and known batches.

The compositional effect size (CES) quantifies batch-induced abundance differences as the mean across glycans of the range (*b*_max_ − *b*_min_) of batch-specific mean CLR abundances.

PCA batch effect is measured as a weighted sum across the first K principal components:

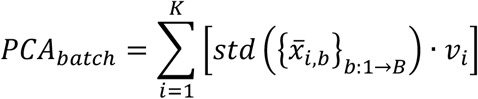

Where *x*_*i,b*_ is the mean *PC*_*i*_ coordinate of samples in batch *b, v*_*i*_ is the explained variance ratio of *PC*_*i*_ and *B* is the number of batches. Higher values indicate stronger batch separation in PCA space.

All distance-based metrics use Euclidean distance on CLR-transformed data to respect compositional geometry.

### Batch effect diagnostic

A rapid diagnostic implemented in the *glycoforge*.*check_batch_effect* function provides immediate assessment of batch effect severity. The function first performs PCA on the CLR-transformed data and extracts principal component scores. When biological group labels are provided, Principal Variance Component Analysis (PVCA) serves as the primary diagnostic metric, computing variance decomposition across the first 10 principal components. Severity classification is assigned based on PVCA batch variance percentage: if below 5%, severity is NONE (negligible batch effect). If batch variance is less than biological variance, severity is GOOD (batch variance below 10%) or MILD (batch variance between 10-20%), indicating biological signal dominates. If batch variance exceeds biological variance, severity escalates based on absolute magnitude: WARNING (batch variance below 20%), MODERATE (20-30%), or CRITICAL (batch variance above 30%). This PVCA-based classification captures batch effects across multiple principal components rather than relying solely on a single axis. Additionally, PC1 analysis provides a complementary single-axis perspective. Statistical testing evaluates whether PC1 scores differ significantly across batches and biological groups. For small sample sizes (n less than 30) or when Shapiro-Wilk normality testing rejects normality at p < 0.05, the nonparametric Kruskal-Wallis test is used; otherwise, one-way ANOVA is applied. Effect sizes are quantified using eta-squared (η^2^), computed as the ratio of between-group variance to total variance for ANOVA or estimated from the Kruskal-Wallis statistic as

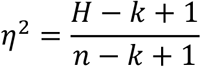

for *k* groups and *n* samples. These PC1 metrics serve as confirmatory evidence alongside the primary PVCA assessment. Finally, the median variance explained by the batch factor across all features is computed by correlating each feature with batch dummy variables and taking the squared maximum correlation per feature, providing a feature-level metric that complements the PC-based global assessment.

### Biological signal preservation metrics

We use two metrics to assess whether batch correction preserves biological structure. The preservation of biological variability metric measures the correlation between F-statistics before and after correction. For each feature, an F-statistic is computed from a one-way ANOVA comparing biological groups, yielding vectors *F*_before_ and *F*_after_. The Pearson correlation between these vectors quantifies how well the feature-wise biological signal strength is maintained, with values closer to 1 indicating better preservation. The conserved differential relationships metric evaluates whether pairwise feature correlations are maintained. Correlation matrices *R*_before_ and *R*_after_ are computed from the data before and after correction. Relationships where the absolute correlation exceeds a threshold *τ* = 0.05 are identified, and the proportion of before-correction relationships that remain above threshold after correction is calculated. This metric captures whether the compositional co-variation structure driven by biological differences survives the batch correction process.

## Data Availability

All data used in this work are available at https://github.com/BojarLab/glycowork.

## Code Availability

The code developed and used in this work can be found at https://github.com/BojarLab/GlycoForge.

## Acknowledgment

This work was funded by a Branco Weiss Fellowship – Society in Science awarded to D.B., by the Knut and Alice Wallenberg Foundation, and the University of Gothenburg, Sweden. The funders had no role in study design, data collection and analysis, decision to publish or preparation of the manuscript.

## Contributions

Conceptualization: D.B., S.H., Funding Acquisition: D.B., Resources: D.B., Software: D.B., S.H., Supervision: D.B., Visualization: D.B., S.H., Writing—Original Draft Preparation: D.B., S.H., Writing—Review & Editing: D.B., S.H.

