## Supplemental Figures for "GlycoForge generates realistic glycomics data under known ground truth for rigorous method benchmarking"

### Supplementary Figures

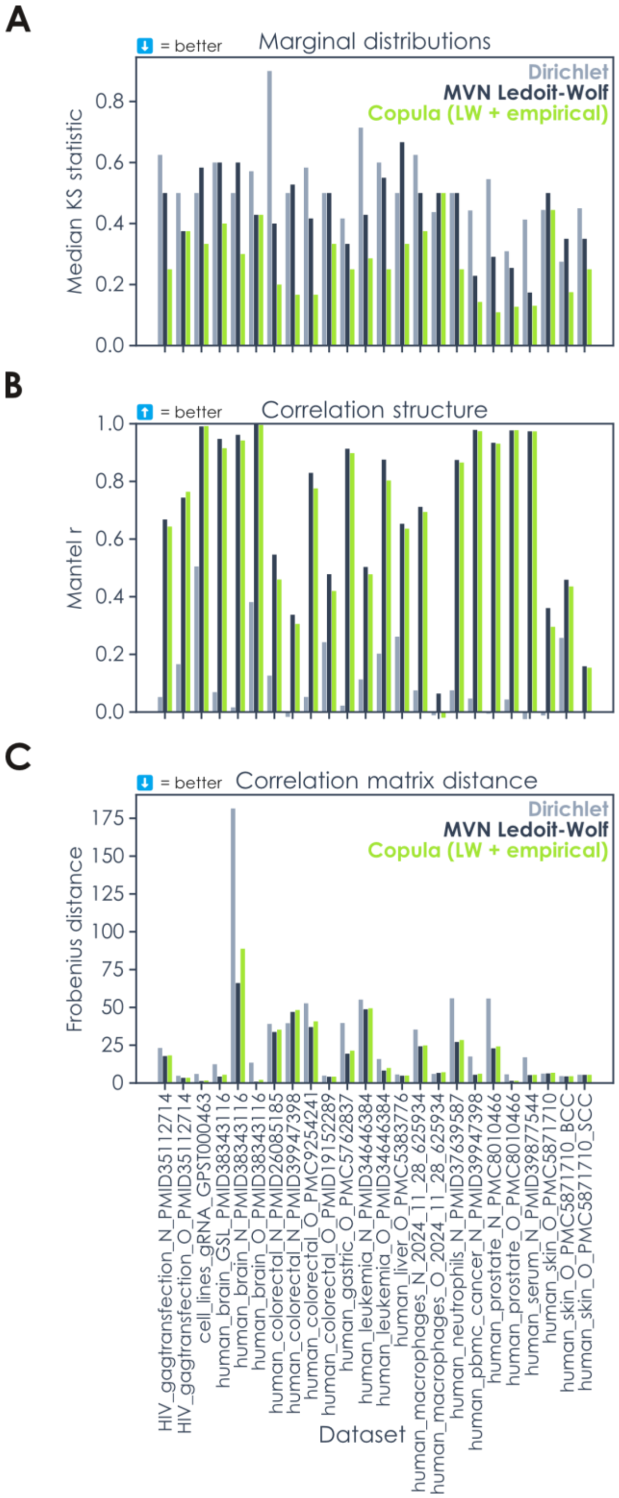


**Figure S1. Comparing simulation mechanisms for glycomics data. A-C)** For all curated glycomics datasets stored in glycowork (v1.8.0) that contained class labels (n = 24), we performed templated simulations via either Dirichlet sampling, multivariate normal sampling with Ledoit-Wolf shrinkage covariance (MVN Ledoit-Wolf), or a Gaussian copula with Ledoit-Wolf and empirical marginals derived from the input dataset. We evaluated goodness-of-fit between simulated and real data via low Kolmogorov-Smirnov statistic (marginal distribution; A), high Mantel r (correlation structure; B), and low Frobenius distance (correlation matrix distance; C).

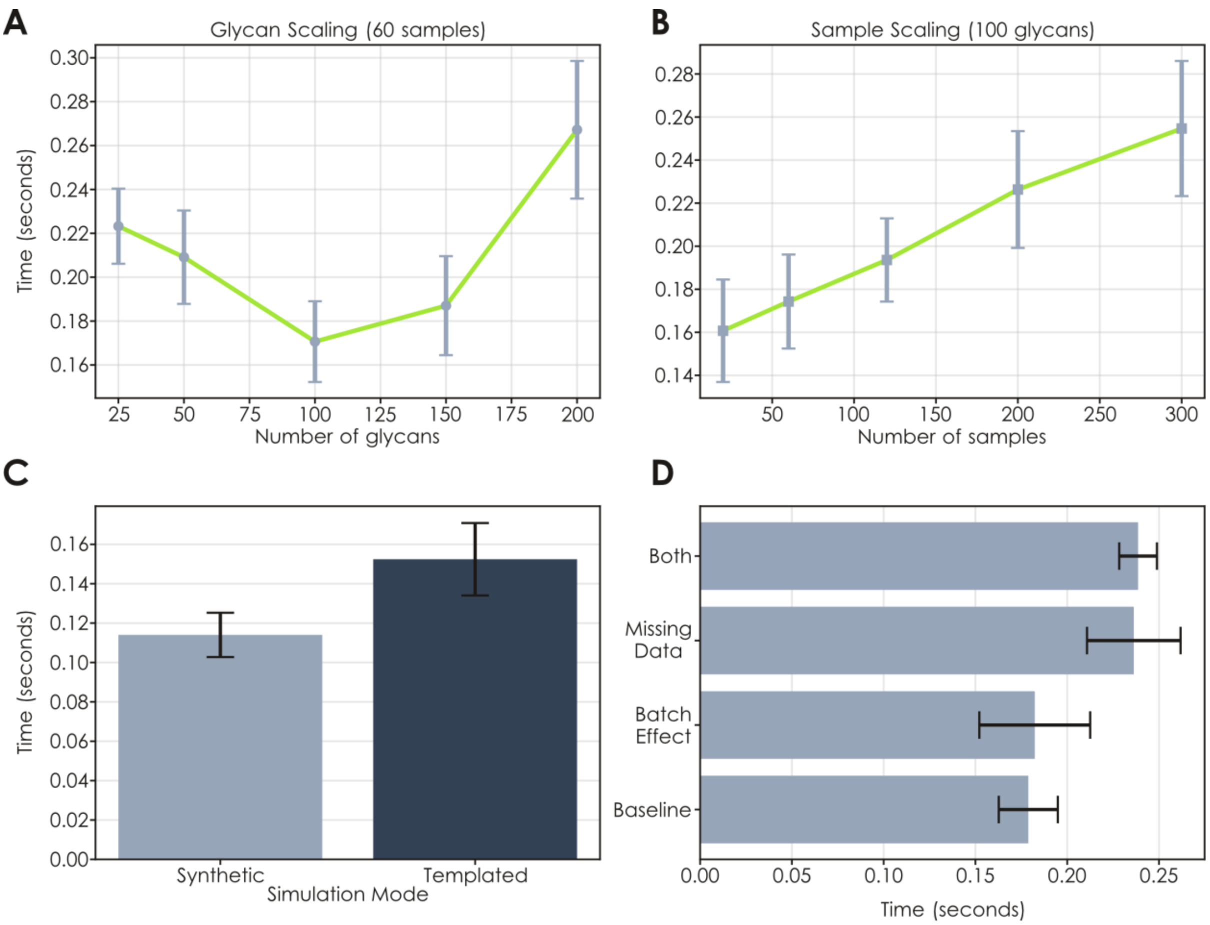


**Figure S2. GlycoForge can produce realistic glycomics data at scale. A-B)** Large glycomics datasets can be simulated in less than one second. Varying either number of glycans (A) or samples (B) maintains scaling compatible with high-throughput simulations. Simulated data configurations (n = 20 seeds per condition) are represented by points, with the data being connected via a line plot. **C)** Timing comparison between synthetic and templated simulation modes. As a template, leukemia *O*-glycomics data (Oliveira et al., *Theranostics*, 2021) were chosen. Settings for both modes included 30 samples per condition and 98 glycans. **D)** Timing additions of including data imperfections in simulation workflow. Missing data insertion used a target fraction of 15%, while the batch effect was set at $\kappa_\mu$ = 1.5 and $var_b$ = 0.5. Settings for all modes included 30 samples per condition and 98 glycans. All data are presented as mean values +/- standard deviation as error bars, stemming from simulations of three independent seeds. All timings were done on a Windows laptop using an Intel® Core™ Ultra 9 185H (2.50 GHz) CPU.

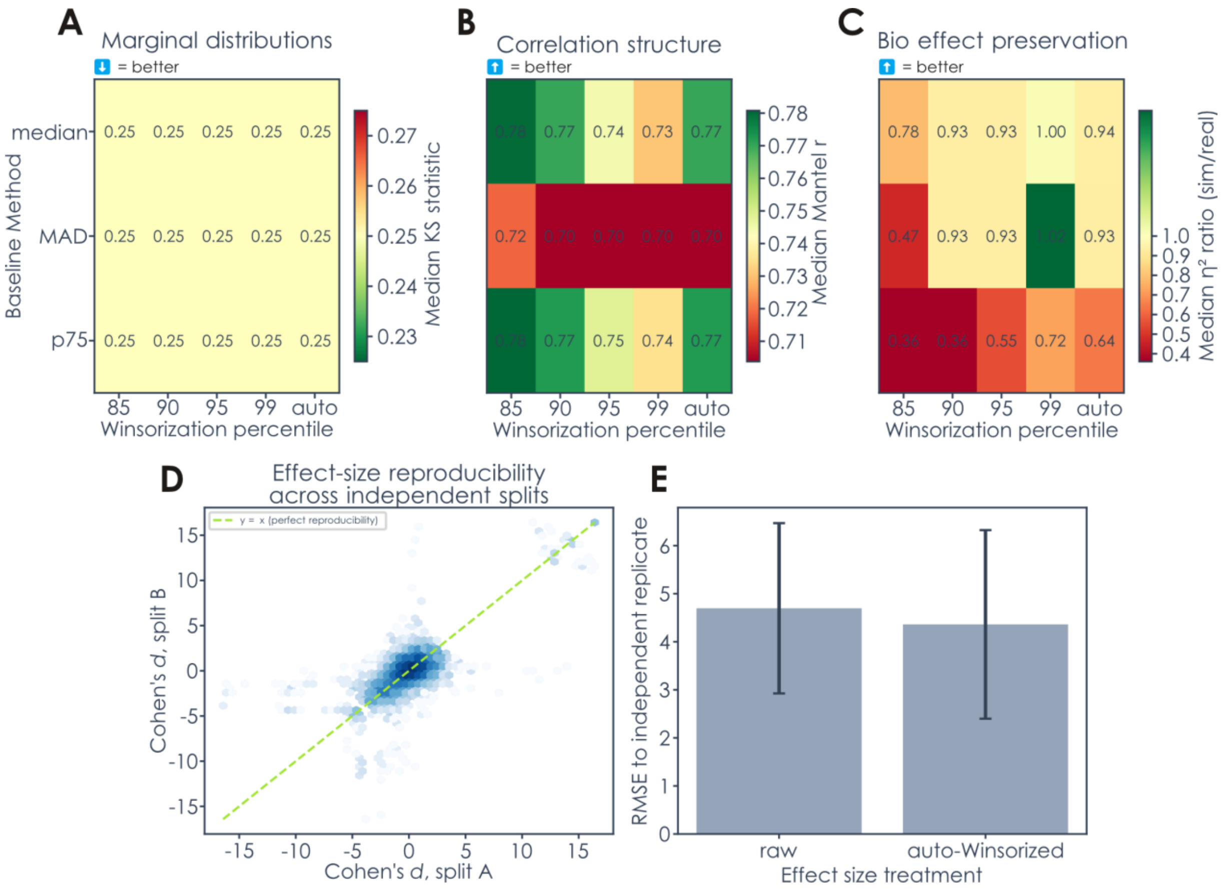


**Figure S3. Sensitivity of templated simulation to Winsorization and baseline normalization parameters. A-C)** Heatmaps show median performance across all 45 glycomics reference datasets stored in glycowork (v1.8.0) as a function of Winsorization percentile (columns) and baseline normalization method (rows), evaluated using Gaussian copula sampling at $bio_{strength}$ = 1.5, $k_{dir}$ = 100, and $variance_{ratio}$ = 1.5. **(A)** Marginal fidelity measured by median two-sample KS statistic between simulated and real CLR distributions (lower is better). All parameter combinations achieve near-identical KS ≈ 0.25, indicating that Winsorization and baseline methods do not affect marginal realism, which is governed entirely by the copula's empirical quantile transform. **(B)** Correlation structure preservation measured by Mantel r between simulated and real pairwise correlation matrices (higher is better). Performance is stable across most combinations (Mantel r ≈ 0.73–0.77) but degrades slightly at too permissive Winsorization and with the MAD baseline method. **(C)** Biological effect preservation measured by the ratio of simulated to real $\eta^2$ on PC1 (optimal at 1.0). The p75 baseline consistently underestimates effect magnitude ($\eta^2$ ratio ≈ 0.44–0.65), while median and MAD baselines achieve ratios closer to unity (0.93-1.00 for most Winsorization thresholds), with the median baseline at the $95^{th}$-$99^{th}$ percentile and auto settings providing the most faithful reproduction of the original biological signal. **(D-E)** Assessing effect-size clipping. For each of the nine benchmark datasets with at least six samples per condition, samples were repeatedly partitioned (50 random splits) into two disjoint, condition-balanced halves, and Cohen's *d* was computed per glycan independently on each half in CLR space. (D) Per-glycan effect sizes from the two independent halves, pooled across all splits and datasets; the dashed line marks perfect reproducibility (y = x). (E) Root-mean-square error between the split-A effect size and its independent split-B replicate (error bars: 95% bootstrap CI), before (raw) and after applying GlycoForge's data-adaptive auto-Winsorization to the split-A estimate, computed over the clipped extreme tail ($|d| > 5$).

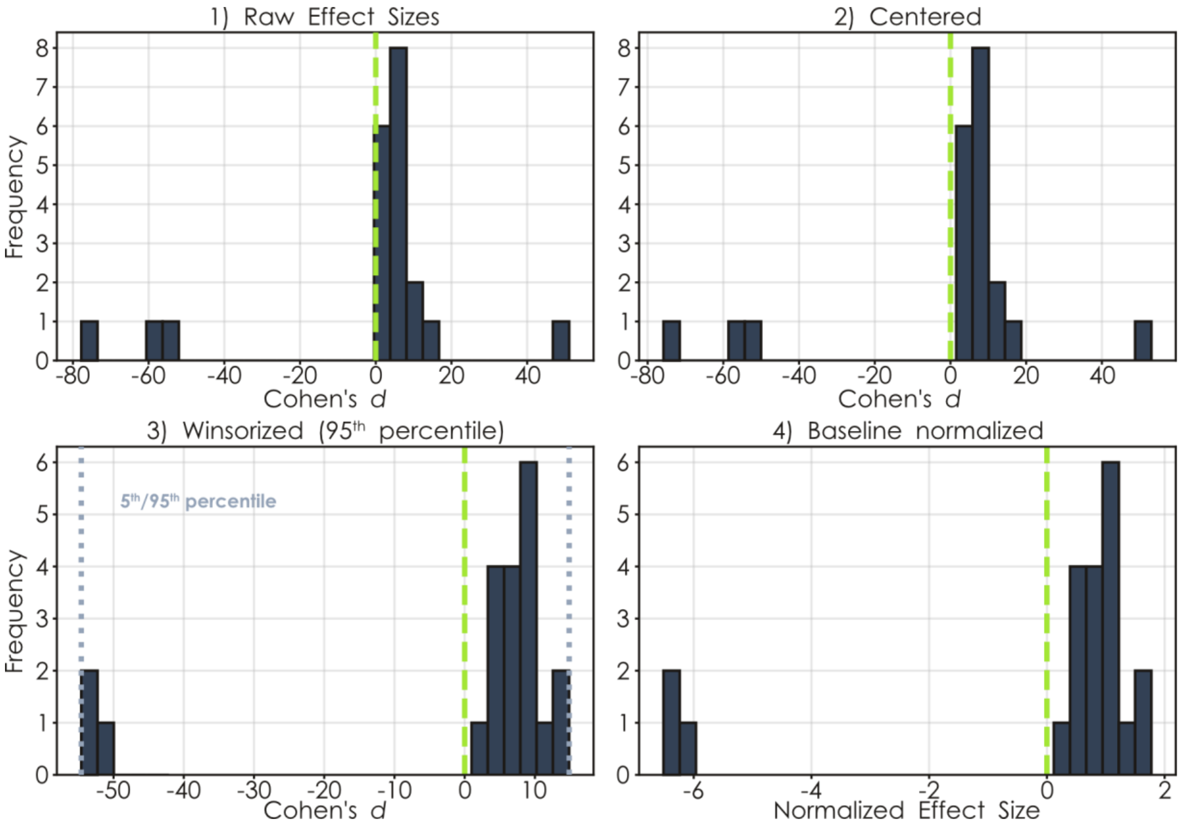


**Figure S4. Showcasing the effect size processing prior to injection in the templated mode.** Raw effect sizes from a leukemia *O*-glycomics dataset (Oliveira et al., *Theranostics*, 2021) were further processed by subtracting the mean, capping outliers via Winsorization, and normalizing effect sizes by dividing them by the median effect size. Data are shown as bar graphs, with visual guides for zero effect size as well as the $5^{th}$ and $95^{th}$ percentile.

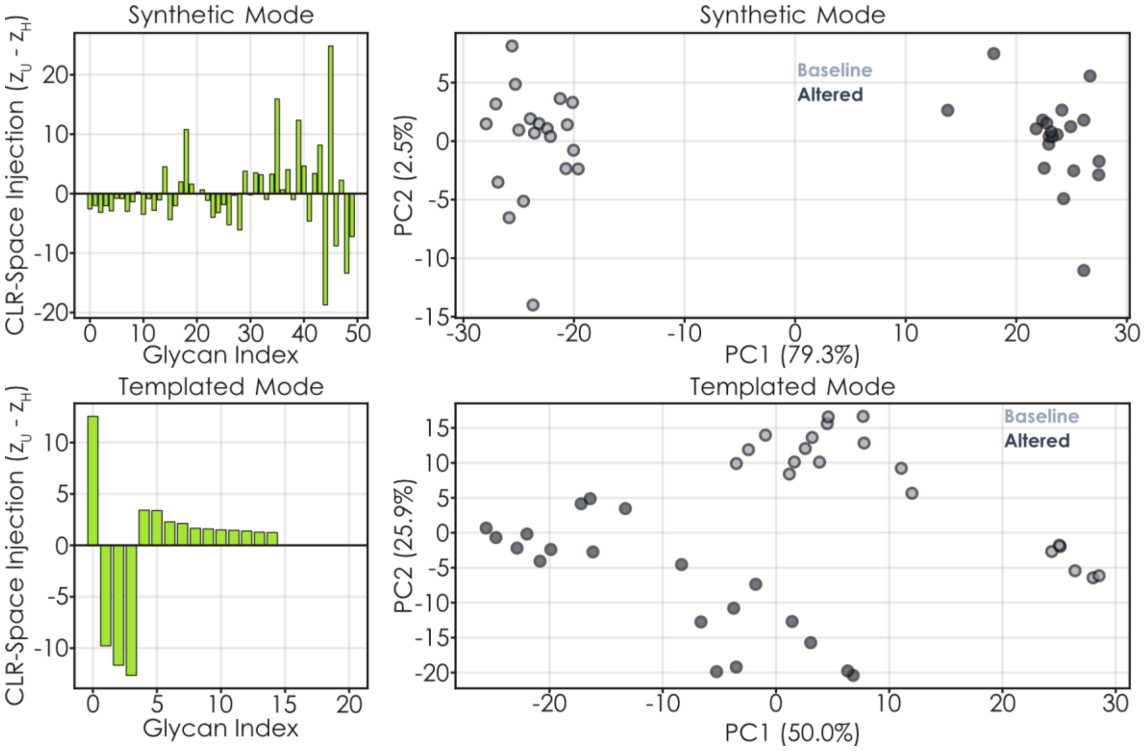


**Figure S5. Comparing synthetic and templated glycomics data generation.** Shown are CLR-space injection vectors as bar graphs, as well as generated glycomes via PCAs. For synthetic, 30% upregulated and 35% downregulated glycans were randomly sampled for the altered cohort. For templated, we used a human GAG-transfected *N*-glycomics dataset (Lavado-Garcia et al., *Biotechnol Bioeng*, 2022) as a template.

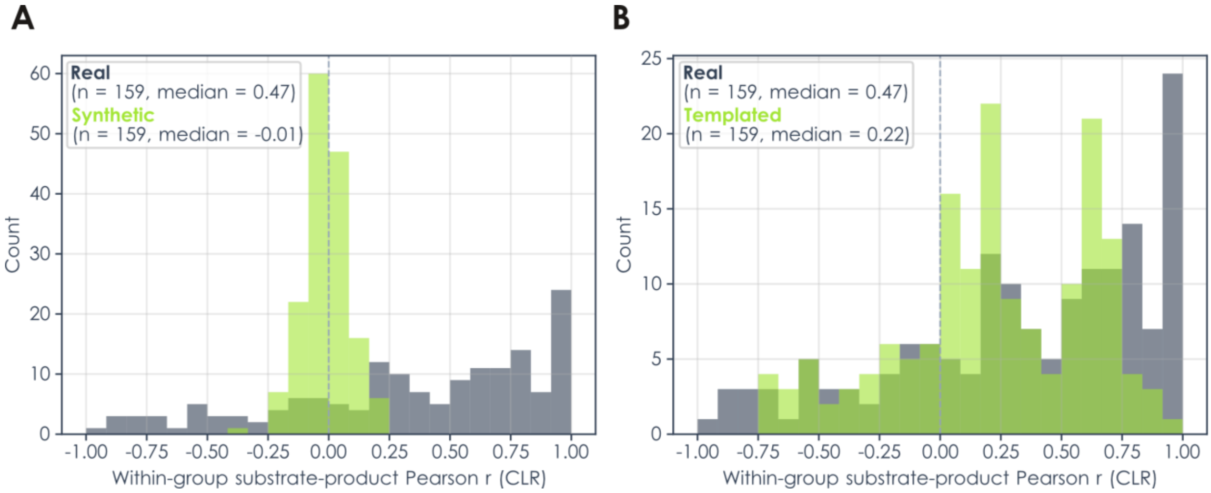


**Figure S6. Within-group substrate-product correlations in real versus simulated glycomics data. A-B)** Distributions of within-group Pearson correlations (CLR space) between Neu5Ac-linked substrate-product pairs, computed separately within healthy and unhealthy groups and averaged, across 13 dysregulated datasets (|mean Neu5Ac effect| ≥ 0.3) from glycowork (v1.8.0). (A) Synthetic mode (Gaussian copula with generic cross-dataset covariance pooled by abundance rank, without glycan-identity alignment) produces near-zero correlations (median = -0.01), reflecting a covariance structure that captures realistic eigenspectrum shape but does not encode biosynthetic relationships between specific glycan positions. (B) Templated mode (Gaussian copula with dataset-specific Ledoit-Wolf covariance and empirical marginals) recovers positive substrate-product correlations (median = 0.22) but underestimates the real data distribution (median = 0.47), consistent with the expected attenuation from pooling covariance across biological groups and the low-n/high-p regime limiting recoverable within-group covariance structure. Grey histograms show real data in both panels; n = 159 substrate-product pairs from the same set of dysregulated datasets are used across all conditions.

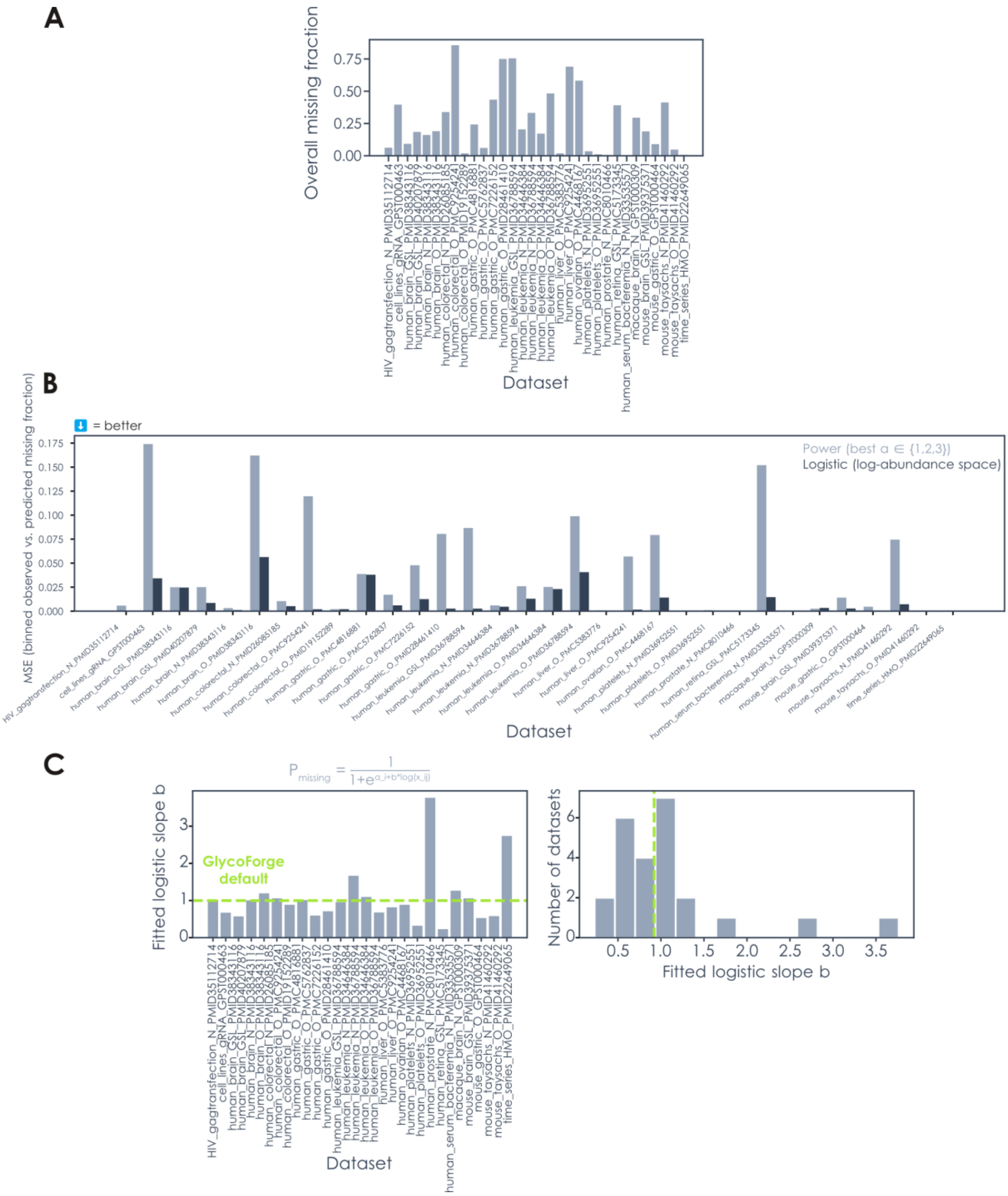


**Figure S7. Empirical validation of an MNAR missingness model across glycowork datasets. A)** Overall missing fraction per dataset (treating zeros as missing; n = 24), showing substantial variability across glycomics studies. **B)** Mean squared error (MSE) between binned observed and predicted missing fractions for two candidate models: a power function in abundance space (best α selected from {1, 2, 3}, one free parameter) and a logistic function in log-abundance space (two free parameters, a and b). The logistic model achieves lower or comparable MSE across the majority of datasets (n = 24; datasets from glycowork v1.8.0), supporting its use as the default MNAR mechanism in GlycoForge. **C)** Per-dataset fitted logistic slope b, which controls the strength of the intensity-dependent detection bias (left), and its distribution across all datasets with valid fits ($0 < b \leq 15$; right). The median fitted slope ($b = 0.92$) empirically justifies GlycoForge's default mnar_bias = 1.0. The logistic model parameterizes per-sample missing probability as $p_{missing} = 1 / (1 + \exp(a_i + b \cdot \log(x_{ij})))$, where $a_i$ is a per-sample intercept solved via Brent's method to achieve the user-specified target missing fraction exactly, and b (= mnar_bias) controls how steeply low-abundance glycans are preferentially lost.

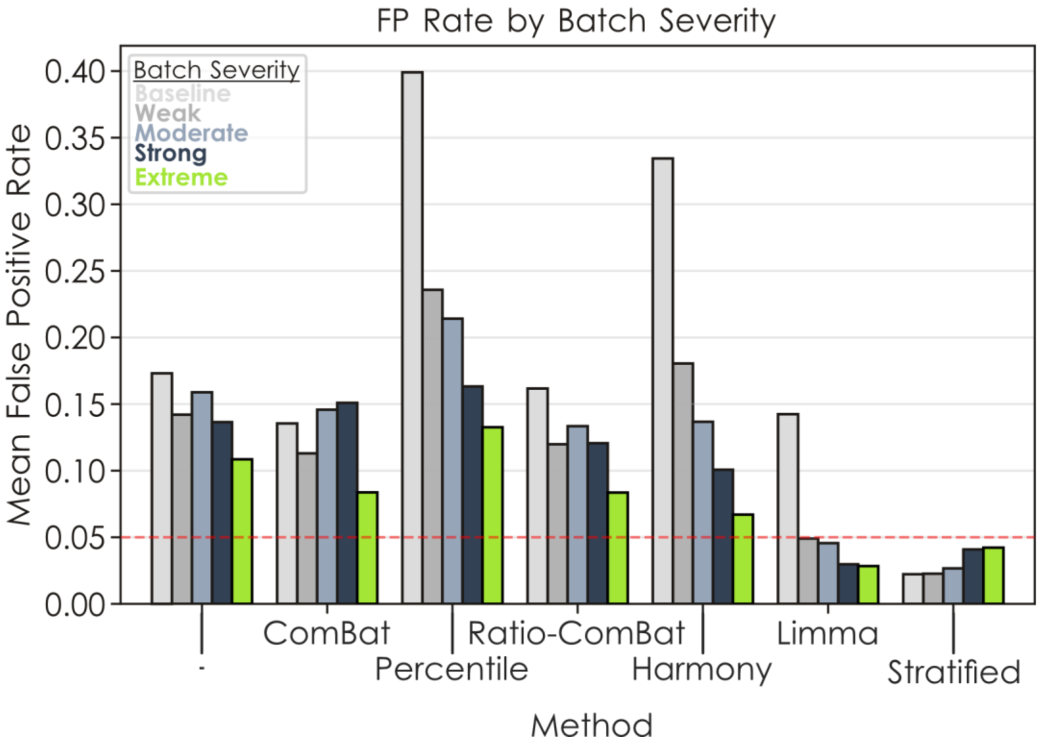


**Figure S8. False-positive rate of batch effect correction methods becomes high for weak batch effects.** Using synthetic glycomics data with increasing batch effect strengths, we assessed the false-positive rate of all tested batch effect correction methods, which is shown as a bar graph. A red dashed line delineates the commonly used and acceptable 5% false-positive rate.

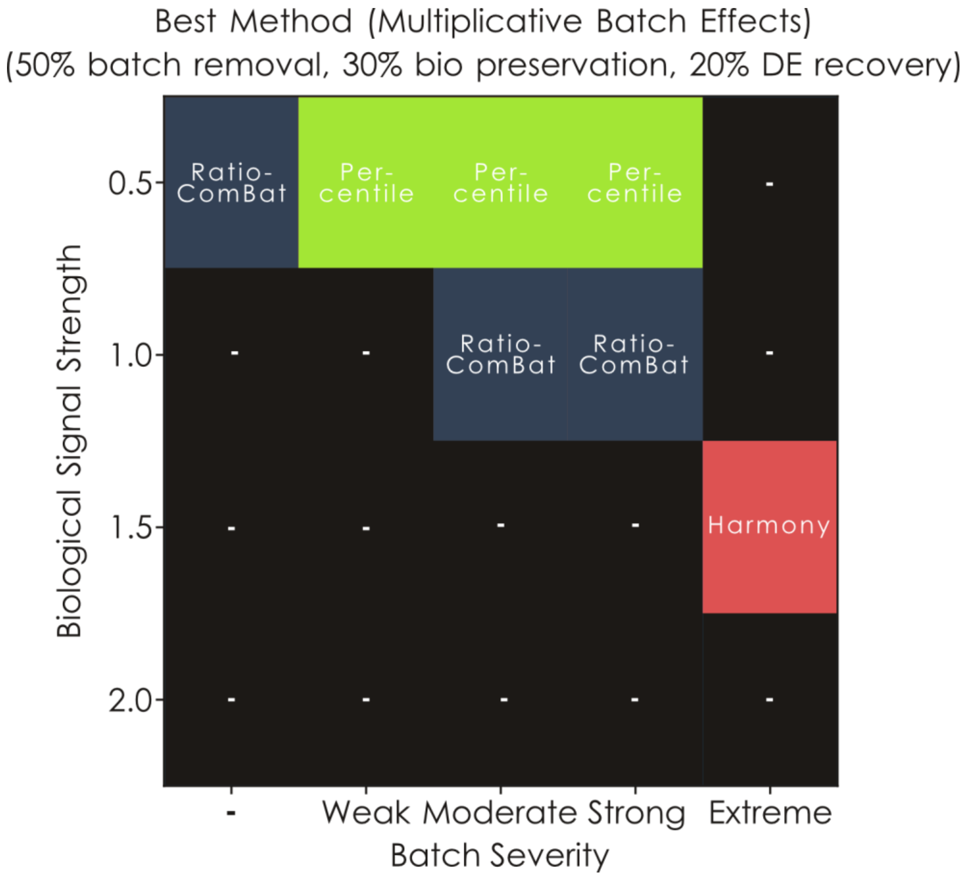


**Figure S9. Benchmarking removal of multiplicative batch effects.** Best-performing batch correction method under multiplicative batch effects, determined by a composite score weighting batch variance removal (50%), biological signal preservation (30%), and differential expression recovery (20%). Batch variance and biological variance were quantified by PVCA decomposition across ten principal components, and DE recovery was measured by F1 score comparing corrected data against the known ground truth from clean simulated data. Each cell reports the winning method for a given combination of biological signal strength ($bio_{\text{strength}} \in \{0.5, 1.0, 1.5, 2.0\}$) and batch severity (sum of mean-shift magnitude $\kappa_\mu$ and variance inflation $var_b$, binned into five levels from Baseline through Extreme). Results were averaged across three random seeds and all parameter combinations within each severity bin. Black cells marked "–" indicate conditions where no correction method outperformed the uncorrected data under the composite criterion. The multiplicative batch effect model applies a log-linear gain factor $\exp(\kappa_\mu \cdot u_b)$ per glycan per batch in CLR space, producing intensity-dependent shifts that scale with each feature's abundance level. Simulations used 50 glycans, 15 healthy and 15 unhealthy samples, and three batches with stratified assignment.

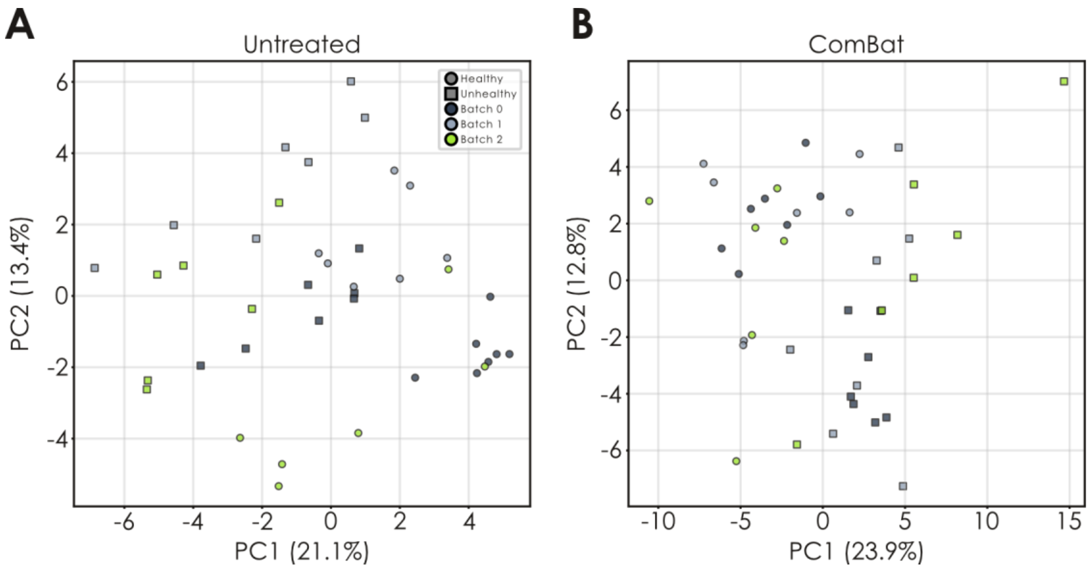


**Figure S10. Removing specific and confounding batch effects from glycomics data. A-B)** Using the glycans from a gastric *O*-glycomics dataset (Adamczyk et al., *Sci Rep*, 2018), we generated synthetic data with a defined decrease in sialylated glycans in the unhealthy group (A). Further, batch effects were injected as a decrease in sialylation in batches 0 and 1. ComBat was able to remove this confounding batch effect (B; PVCA batch variance 27.37% to 0%; kBET 3.9 to 1.7, ARI 0.47 to -0.02). Biological status is indicated by shape, whereas batch is indicated by color.

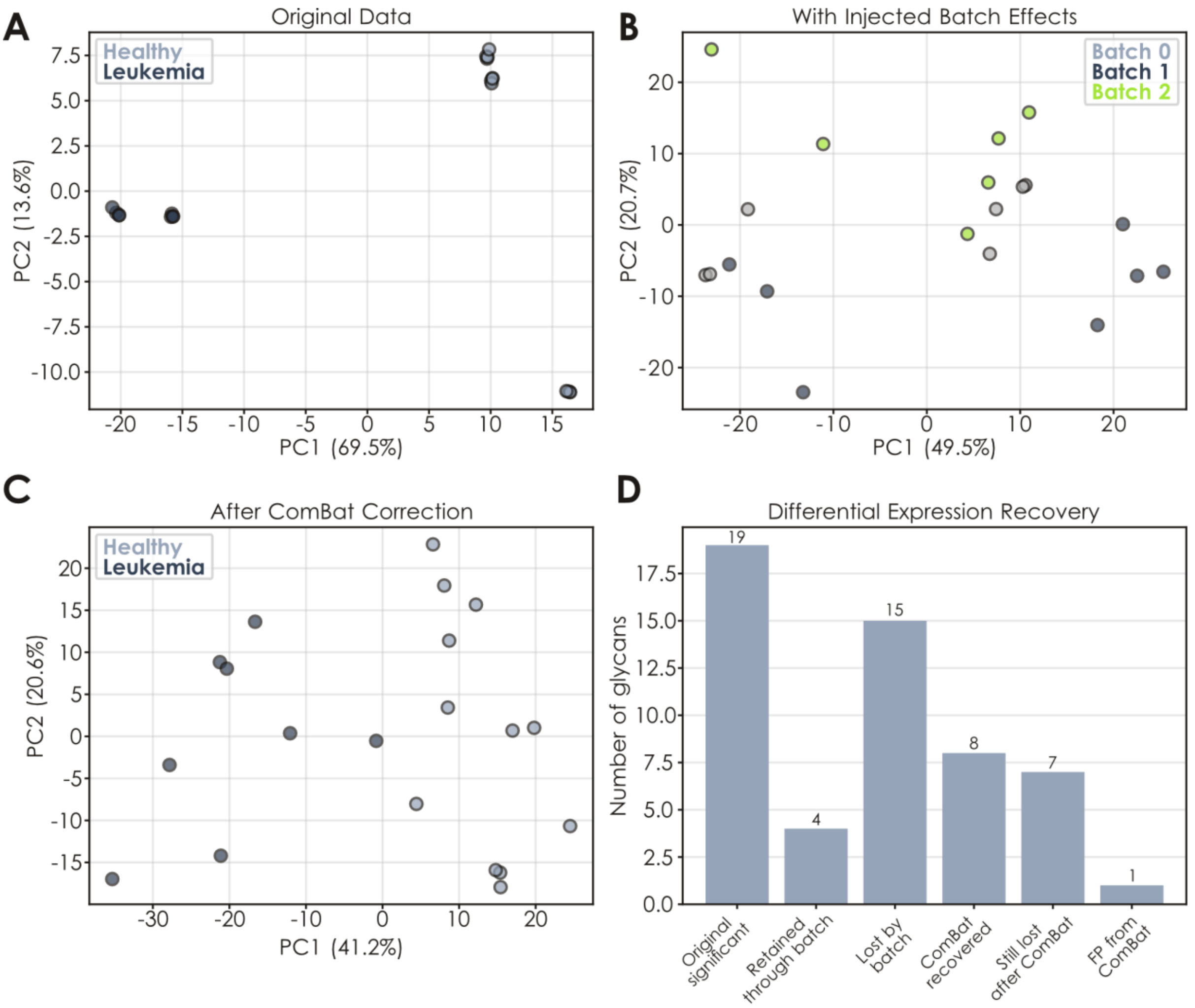


**Figure S11. Batch effects injected into real glycomics data can be removed by ComBat. A-C)** Injecting moderate batch effects ($\kappa_\mu$ = 0.6 and $var_b$ = 0.3) into leukemia *O*-glycomics data (Oliveira et al., *Theranostics*, 2021; A) via GlycoForge (B) that can be corrected via ComBat (C) is shown via PCA plots. Batches are created via random class-stratified sampling. **D)** Assessing ComBat performance on batch effect correction in real-world data. Using the *glycowork.motif.analysis.get_differential_expression* function, we profiled significant differences between healthy and leukemia samples in the original data, the batch effect injected data, and the ComBat-corrected data, visualized via bar graphs, to conclude that ComBat recovers >50% of lost significant effects in this case.

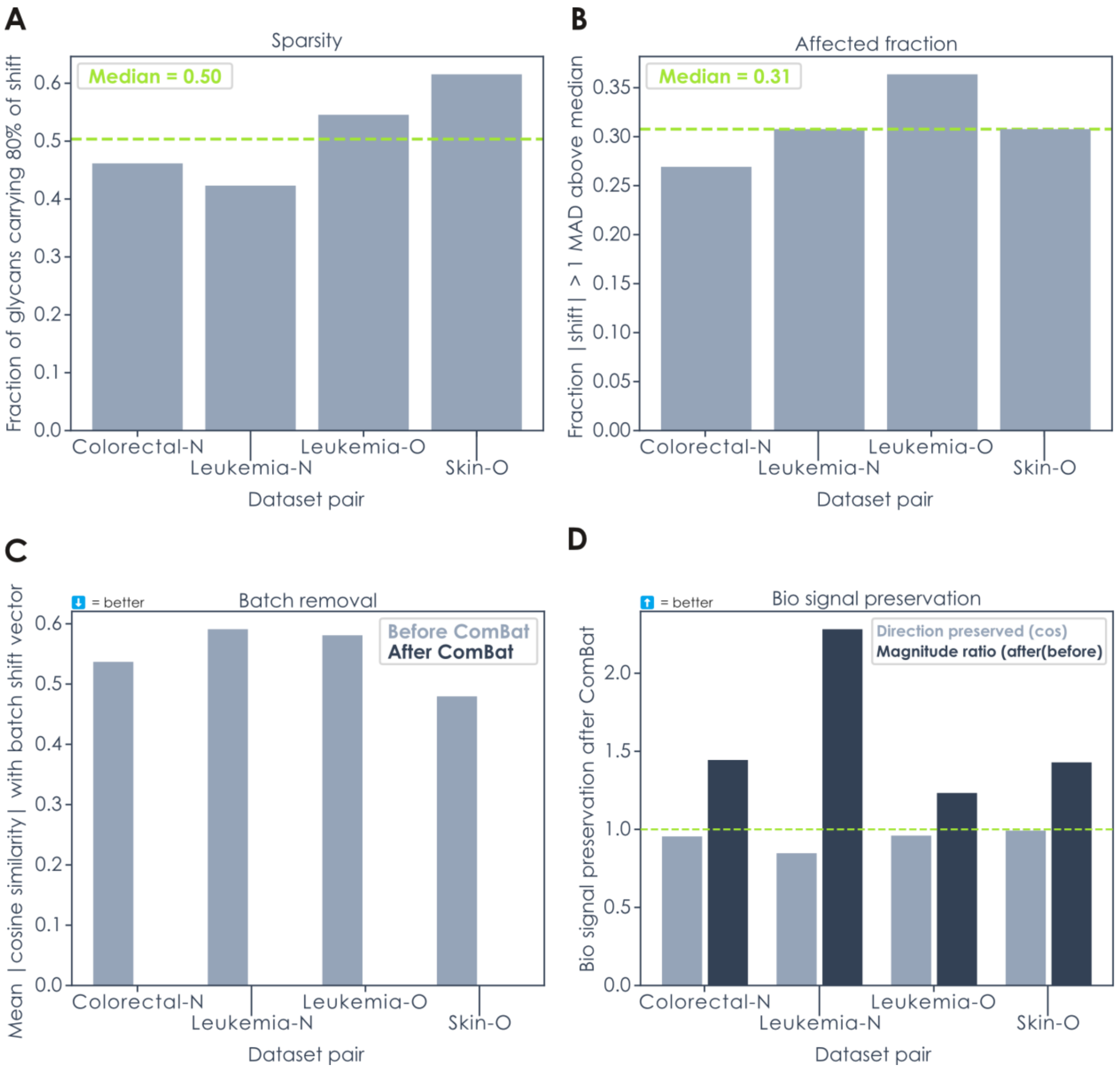


**Figure S12. Cross-study batch effect structure in real glycomics data. A-D)** Four pairs of independently published glycomics datasets sharing the same tissue and glycan class were aligned on common glycan sequences, CLR-transformed, and compared before and after ComBat correction. **A)** Sparsity of the between-study CLR shift, measured as the fraction of glycans carrying 80% of the total shift magnitude (median = 0.50), indicating that batch effects are distributed across roughly half the glycome. **B)** Fraction of glycans whose absolute shift exceeds one MAD above the median (median = 0.31), providing an independent estimate of the affected glycan proportion. **C)** Mean absolute cosine similarity between individual sample deviations and the batch shift vector before and after ComBat; near-complete removal confirms that the observed shifts are systematic and correctable. **D)** Biological signal preservation after ComBat, quantified as the cosine similarity between within-study healthy-versus-unhealthy CLR deltas before and after correction (direction preservation) and the ratio of their L2 norms (magnitude ratio). Direction is fully preserved (median cosine ≈ 0.96), while magnitude increases modestly (median ratio > 1), consistent with reduced noise masking after batch removal.

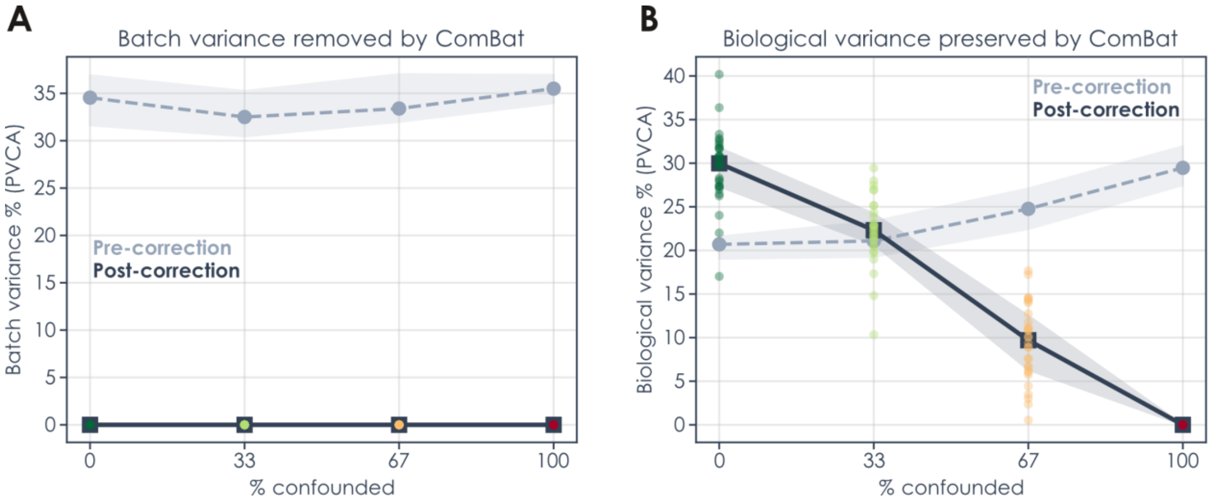


**Figure S13. Batch-biology confounding degrades ComBat's ability to preserve biological signal. A-B)** Simulated glycomics datasets were generated with increasing levels of confounding between batch assignments and biological groups (0% = fully stratified, 100% = fully confounded), and corrected using ComBat. (A) Batch variance (PVCA) is effectively removed across all confounding levels, confirming that ComBat consistently eliminates technical variation regardless of study design. (B) Biological variance retained after correction decreases monotonically with confounding severity, declining from ~30% under stratified assignment to near-zero under full confounding, where batch and biological effects become indistinguishable and ComBat removes both. Lines show median PVCA across simulation runs (dashed gray: pre-correction; solid black: post-correction), shaded regions span the interquartile range, and individual run results are shown as colored points.

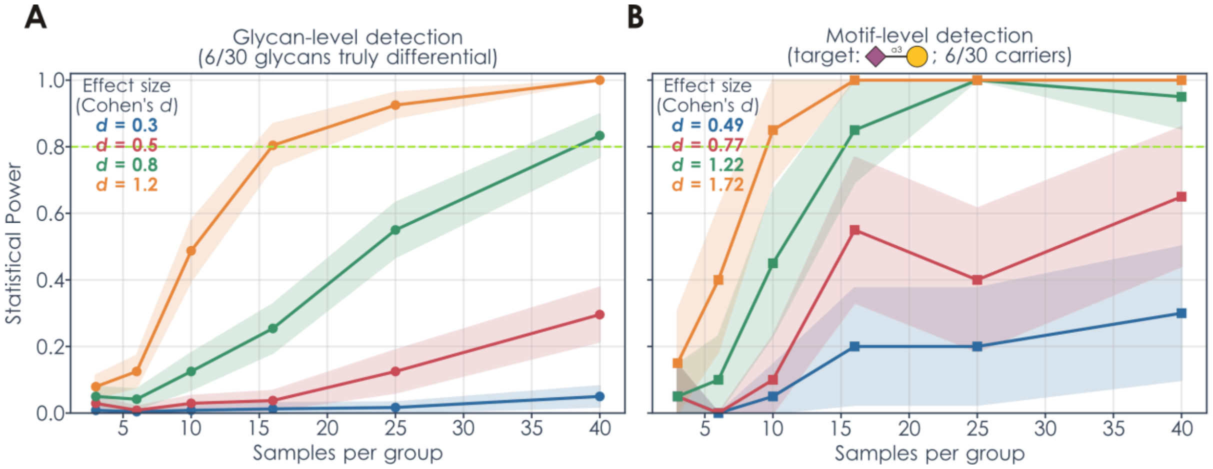


**Figure S14. Statistical power estimations using GlycoForge-simulated glycomics data. A-B)** Synthetic compositional datasets were generated by Gaussian copula sampling using empirical centered-log-ratio (CLR) marginals and a Ledoit-Wolf shrinkage covariance estimated from the 30 most abundant *N*-glycans of the reference dataset (human_serum_bacteremia_N_PMID33535571), yielding two equal-sized groups across the sample sizes shown on the x-axis. (A) Glycan-level detection: 6 of 30 glycans were designated differential and assigned balanced up-/down-regulation injected in CLR space at the nominal Cohen's *d* indicated in the legend. (B) Motif-level detection: the 6 glycans carrying Neu5Ac α2-3Gal were coherently regulated, with legend values giving the resulting motif-level Cohen's *d* (measured from the aggregated motif at the largest sample size). For each condition, differential expression was assessed with glycowork's CLR-based test and Benjamini-Hochberg correction (per-glycan in A, on aggregated motif abundances in B); power is the mean fraction of truly differential features called significant across independent simulations (40 in A, 20 in B). Shaded regions are 95% Monte Carlo confidence intervals across simulations; the dashed line marks 80% power.
